# Strong and efficient biological carbon pump in the Northern Gulf of Alaska during summer

**DOI:** 10.1101/2023.11.10.566633

**Authors:** Stephanie H. O’Daly, Gwenn M. M. Hennon, Thomas B. Kelly, Suzanne L. Strom, Andrew M. P. McDonnell

## Abstract

Sinking marine particles, one pathway of the biological carbon pump, transport carbon to the deep ocean from the ocean’s surface, thereby contributing to atmospheric carbon dioxide modulation and benthic food supply. Few *in situ* measurements exist of sinking particles in the Northern Gulf of Alaska (NGA); therefore, regional carbon flux prediction is poorly constrained. In this study, we aim to (1) characterize the magnitude and efficiency of the biological carbon pump and (2) identify drivers of carbon flux in the NGA. We deployed drifting sediment traps to simultaneously collect bulk carbon and intact sinking particles in polyacrylamide gels and measured net primary productivity from deck-board incubations. Through deployments during the summer of 2019, we found high carbon flux magnitude, low attenuation with depth, and high export efficiency. We quantitatively attributed carbon flux between ten particle types, including various fecal pellet categories, dense detritus, and aggregates using polyacrylamide gels. The contribution of aggregates to total carbon flux (41 - 93%) and total carbon flux variability (95%) suggests that aggregation processes, not zooplankton repackaging, played a dominant role in carbon export during the summer of 2019 in the NGA. Furthermore, efficient export correlated significantly with the proportion of chlA > 20 µm, total aggregate flux, and proportion aggregate flux. These results suggest that this stratified, small-cell-dominated ecosystem can have sufficient aggregation to allow for a strong and efficient biological carbon pump. These are the first measurements of carbon flux and the first integrative description of the BCP in this region.

**Significance Statement:** *Novelty and significance:* We use a comprehensive approach that brings together sediment trap sampling and imaging, optically measured distribution of sinking and suspended particles, and incubations to make the first description of the biological carbon pump in the Northern Gulf of Alaska. We found high carbon flux magnitude, low attenuation with depth, and high export efficiency with a phytoplankton community consisting of mostly pico-and nanoplankton. Notably, just 25% of carbon flux out of the euphotic zone was as recognizable fecal pellets; instead, we demonstrate that aggregation processes were the main driver of carbon flux. Additionally, size-fractionated chlorophyll-*a* (> 20 µm) strongly correlated with export efficiency across our region. These results lead us to question our expectations about what conditions and processes can create strong and efficient flux events in the Gulf of Alaska.

*Breadth of Interest:* This study is the first description of the biological carbon pump in the Northern Gulf of Alaska and greatly improves biogeochemical constraints on this system. We report observed primary production, carbon flux, export ratio, carbon flux attenuation, and carbon flux by 10 particle types, which can be used to test regional climate models. This study builds on previous studies published in L&O: Strom et al. 2007; Ebersbach & Trull 2008; McDonnell & Buesseler 2010, 2012; and Durkin et al. 2016.

**Author contribution statement:** SO, SS, and AM: conceptualization, methodology, and investigation. SS, AM, GH: funding acquisition and project administration. SO, TK, AM: formal analysis. GH, TK, and AM: supervision. SO: visualization, writing-original draft preparation. SO, GH, TK, SS, and AM: writing-reviewing and editing.

## Introduction

Gravitationally sinking marine particles transport photosynthetically fixed carbon from the euphotic zone into the ocean’s interior and benthos. Such flux is an important biological carbon pump (BCP) component, contributing to the global carbon cycle (Kwon et al. 2009; le Moigne 2019). Despite many decades of studies, there are large uncertainties in our understanding of the BCP due partly to large temporal and spatial heterogeneity in export flux (Buesseler et al. 2007; Henson et al. 2011; Laufkötter et al. 2016; Kelly et al. 2018). While global BCP estimates vary by nearly 50%, ranging between 7 – 16 Pg C year^-1^ (Dunne et al. 2007; Henson et al. 2012; Laufkötter et al. 2016), regional estimates under a changing climate are often even more uncertain. For example, some models (Weber et al. 2016) predict more efficient carbon export in high-latitude regions, such as the Northern Gulf of Alaska (NGA), under warming ocean conditions while others predict less efficient carbon export (Henson et al. 2012; Marsay et al. 2015).

The composition of sinking marine particles affects the overall net carbon flux (Turner 2015); therefore, accurately parameterizing rates based on this composition is an important way to improve climate model accuracy (Siegel et al. 2014; Laufkötter et al. 2016). Marine particles can be broadly organized into three functional types: (1) phytoplankton cells, (2) zooplankton fecal pellets, and (3) aggregates of detrital material (Turner 2015). These different types of particles overlap in size but distinct ranges in composition, density, shape, porosity, and ecological sources (Buesseler et al. 2007; Turner 2015), which result in different sinking speeds (Armstrong et al. 2002; De La Rocha and Passow 2007; Iversen and Lampitt 2020) and remineralization rates (Ploug et al. 2008b; Kobari et al. 2013). Marine particle size has been used to calculate carbon content and to predict sinking speed (Stemmann et al. 2004; Guidi et al. 2008); however, considering only particle size masks potential differences in export between particle types. Studying particle types, in addition to size, will help reduce uncertainty in the ocean’s BCP.

While the BCP is poorly studied in the NGA, primary productivity and the phytoplankton community composition have been well studied in this region. Summers are often characterized by a nutrient-limited community of small-celled nanoflagellates and picoplankton (Strom et al. 2006). Stations closer to the coast are often nitrate-limited while stations on the slope are likely iron-limited (Aguilar-Islas et al. 2016). This results in moderate primary productivity rates, similar to other nutrient-limited regions. In contrast, the springtime community is typically characterized by large diatoms and high rates of primary productivity (Strom et al. 2016). Based on the phytoplankton community composition, high rates of microzooplankton grazing (Strom et al. 2001, 2007) and high macrozooplankton biomasses (Moriarty et al. 2013), we would expect a weak and inefficient summertime BCP with sinking material dominated by zooplankton fecal pellets. Indeed, relatively high contributions of fecal pellets have been measured in the Southern Ocean, another high-latitude system (Ebersbach and Trull 2008; Laurenceau-Cornec et al. 2015).

As a part of the NGA Long Term Ecological Research (NGA LTER) program, we used PIT-style (Knauer et al. 1979) sediment traps to measure the quantity and quality of sinking particles in the NGA during June and July 2019, along with corresponding measurements of euphotic zone primary production. Additionally, we estimated the contribution of phytoplankton cells, aggregates, and fecal pellets to overall carbon flux using polyacrylamide gel traps. Our goal was to characterize the strength and efficiency of the BCP across the NGA and the potential drivers of these patterns.

## Materials and Methods

### Study area and hydrography

Data from this study were collected as part of the Northern Gulf of Alaska Long Term Ecological Research (NGA LTER) program during the SKQ2019-15S cruise from June 30 until July 17 of 2019 on the R/V *Sikuliaq*. Three transects were sampled including the Gulf of Alaska line (GAK), the Kodiak Island line (KOD), and the Middleton Island line (MID), as well as stations in Prince William Sound (PWS) (Fig. 1, Table 1).

**Fig. 1.**
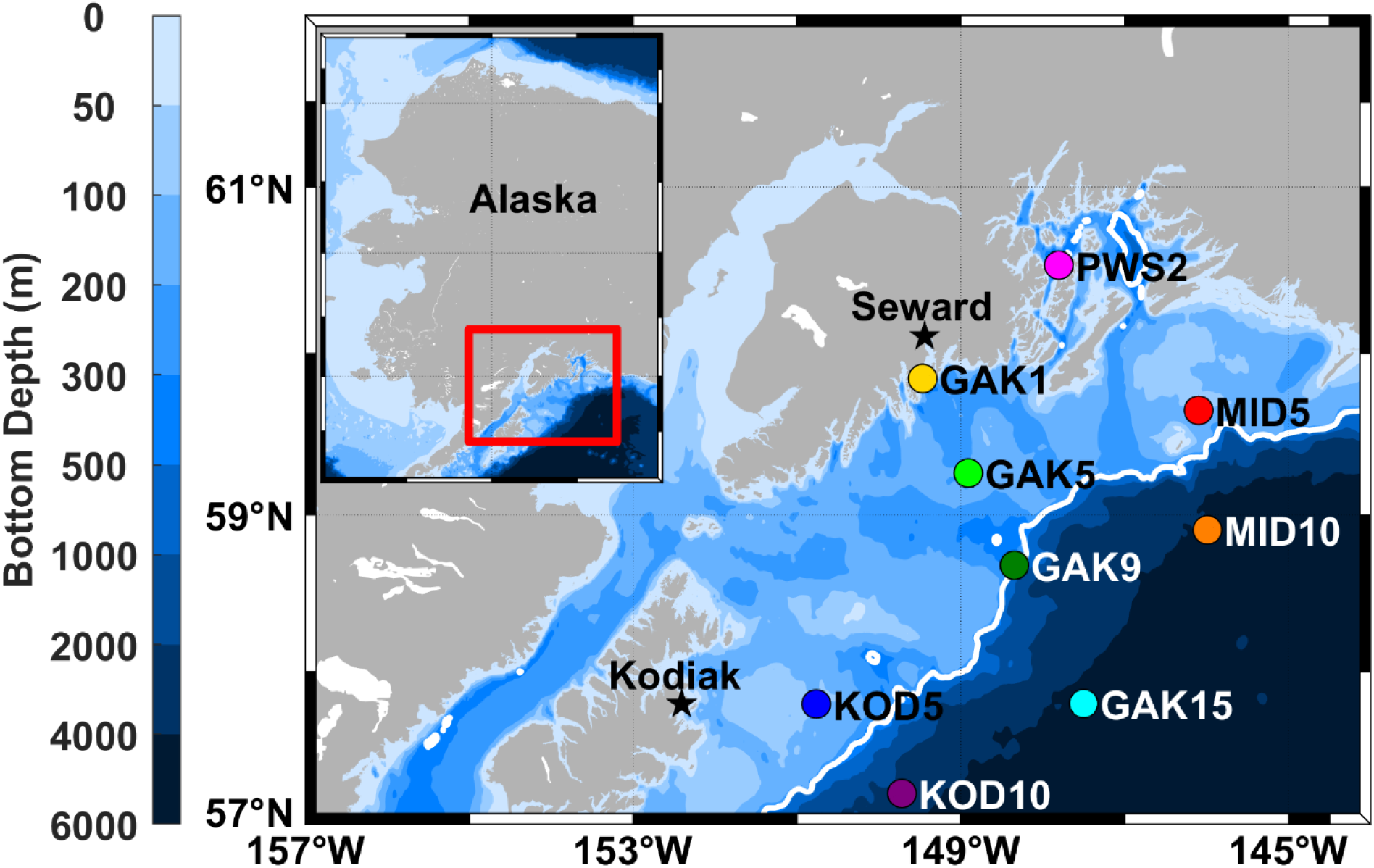
Locations of process stations where drifting sediment traps were deployed and net primary productivity (NPP) rates were measured (colored circles). The 300 m isobath, which separates the continental shelf from slope, is shown in white. These stations largely fall on three transect lines, Kodiak Line (KOD), Gulf of Alaska Line (GAK), and Middleton Island Line (MID), as well as in Prince William Sound (PWS).

**Table 1.** Location, duration, and depth of drifting sediment trap deployments in the Northern Gulf of Alaska (NGA).

| Station | Category | Seafloor depth (m) | Depth of traps (m) | Deploy time (UTC) | Deploy latitude | Deploy longitude | Recover time (UTC) | Recover latitude | Recover longitude | Duration (h) |
| --- | --- | --- | --- | --- | --- | --- | --- | --- | --- | --- |
| GAK1 | Shelf | 269 | 27, 53, 78, 103, 128 | 7/13/19 13:00 | 59.829 | -149.453 | 7/13/19 20:50 | 59.816 | -149.466 | 7.8 |
| GAK5 | Shelf | 173 | 27, 53, 78, 103 | 7/12/19 12:47 | 59.245 | -148.898 | 7/12/19 20:18 | 59.254 | -148.956 | 8.5 |
| GAK9 | Shelf | 270 | 27, 53, 78, 103, 128 | 7/10/19 14:07 | 58.662 | -148.342 | 7/11/19 7:05 | 58.645 | -148.350 | 17.0 |
| GAK15 | Slope | 4400 | 27, 53, 78, 103, 128 | 7/9/19 6:32 | 57.780 | -147.476 | 7/10/19 0:55 | 57.800 | -147.455 | 18.4 |
| KOD5 | Shelf | 86 | 27, 53 | 7/15/19 14:58 | 57.780 | -150.760 | 7/15/19 21:19 | 57.789 | -150.899 | 6.3 |
| KOD10 | Slope | 2501 | 27, 53, 78, 103, 128 | 7/17/19 0:21 | 57.203 | -149.721 | 7/17/19 15:54 | 57.084 | -149.738 | 15.5 |
| MID5 | Shelf | 93 | 25, 50 | 7/2/19 7:31 | 59.65 | -146.095 | 7/2/19 22:11 | 59.652 | -146.168 | 14.7 |
| MID10 | Slope | 4464 | 25, 50, 75, 100, 125 | 7/3/19 13:26 | 58.909 | -145.997 | 7/3/19 21:17 | 58.849 | -145.849 | 7.9 |
| PWS2 | Shelf | 731 | 80, 105, 130, 155 | 6/30/19 17:31 | 60.554 | -147.793 | 7/1/19 3:39 | 60.558 | -147.769 | 10.1 |

Sediment trap, ^13^C net primary productivity (NPP), and CTD data from this study come from process stations (Fig. 1). The CTD unit consisted of a Seabird SBE16plus unit coupled with a WetLabs fluorometer and transmissometer. An Underwater Vision Profiler 5 Standard Definition (UVP5; Hydroptic) unit was mounted to the frame, measured in situ particles from 102 µm to 26 mm, and was deployed with every CTD cast.

### Sediment trap sampling and analysis

Lagrangian surface-tethered drifting sediment traps (PITs; KC Denmark model number 28.2000) were deployed to collect sinking particles (Moran et al. 2012; O’Daly et al. 2020). Nine stations were sampled with durations between 6 - 19 h (median: ∼10 h) depending on other cruise operations (Table 1). The shallowest crossframe was deployed at approximately 27 m, our best estimate of the base of the euphotic zone (1% light level) in this region during the summer based on previous observations, with each subsequent crossframe tethered in 25 m increments. We sampled between two and five depths depending on the local bathymetry at each deployment (Table 1). The deepest trap for each deployment was positioned at least 20 m above the seafloor to reduce the impacts of benthic resuspension. The depth of the euphotic zone during this cruise averaged 28 m. For export flux estimates, we selected the trap that was closest in depth to the base of the euphotic zone (± 9 m) or the trap immediately below the euphotic zone.

Each inline crossframe consisted of four tubes on beveled hinges. Two of the four tubes collected sinking particles in bulk and were filled with filtered seawater brine (i.e., 0.22 µm + NaCl, salinity > 50) that was poisoned (i.e., 1% formalin, final concentration, buffered to saturation with borax) and chilled (4˚C). The remaining two tubes in each trap array had a removable clear-bottomed cup filled with clear viscous polyacrylamide gel (50 mL). The cups were fitted with a thin sloping ramp, which created a seal on the inside of the tube walls and funneled sinking particles into the cup. These tubes were then filled with chilled (4˚C) and filtered (0.22 µm millipore sterivex filter cartridges) seawater.

We measured sinking particulate organic carbon (POC), particulate nitrogen (PN), particulate inorganic carbon (PIC), δ^13^C and δ^15^N isotopic composition, and total suspended particulate matter (SPM) from the bulk trap material as follows. After recovery, each trap tube was covered and left to rest for about two hours until all sinking particles reached the bottom, then overlying water was removed from all tubes down to a boundary layer above the settled particles for the bulk tubes or as close to the surface of the gel as possible without disturbing the gel. Gel cups were removed, covered, and stored in the dark until they could be imaged, which occurred < 6 h after recovering the drifting sediment trap (see “Gel trap imaging and image processing” for more details). Sinking particles in the two bulk collection tubes at each depth were quantitatively split into 10 subsamples each using a McLane rotary splitter. All subsamples were passed through a 500 µm mesh to remove swimmers (Owens et al. 2013; Baker et al. 2020). Three subsamples were combined for POC/PN and an additional three for PIC; combined subsamples were filtered onto precombusted, preweighed 25-mm Whatman GF/F filters. All filters were dried in a dehydrator at 60°C for 12 h and sealed in Petri dishes until further analysis. This resulted in two POC flux values per trap depth (i.e., one per bulk collection tube).

Carbon, nitrogen, and isotopic samples were processed in the Alaska Stable Isotope Facility at the University of Alaska Fairbanks Water and Environmental Research Center. POC/PN filters were acidified with 10% hydrochloric acid for 24 h (fumigation method) to remove inorganic carbon. All sample filters were pelletized in tin cups and processed by CHN elemental analyzer. PIC flux was calculated by subtracting the acidified carbon flux from the non-acidified carbon flux. POC values were converted to daily fluxes (i.e., g C m^-2^ d^1^) based on deployment duration and trap area.

An average chemically derived POC flux error (found to be 14%) was calculated as the average normalized variation from the mean of duplicate samples at each depth (Supporting Information Table 1, Supporting Information Fig. 1). Chemically derived POC flux attenuation was calculated using a power-law function shown in Equation 1 (Buesseler et al. 2007)

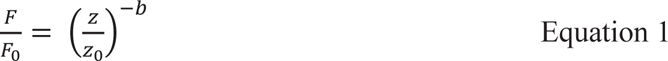

where z is the depth of the trap and z_0_ is the depth of the shallowest trap (m for both), F is the flux at depth z, F_0_ is the flux at the shallowest trap depth, and b is a unitless exponent that is the best-fit attenuation coefficient. The attenuation coefficient, b, was found by fitting Equation 1 and minimizing the sum of squared residuals between measured and modeled POC flux using fminsearch in Matlab 2019B.

### Net primary productivity rate and chlorophyll-*a* measurements

Rates of NPP were measured at 6 depths spanning the euphotic zone using 24 h on-deck incubations (Hama et al. 1983). Water was collected by Niskin rosette, spiked with H^13^CO_3-_ and incubated in screened bags corresponding to *in situ* collection irradiance (range 100 to 1% of surface irradiance as estimated from the PAR attenuation coefficient determined during the downcast). After 24 h, each incubation was filtered onto pre-combusted GF/F filters and frozen at -80°C. Back in the lab, samples were acidified (fumed) with 10% hydrochloric acid for 24 h to remove PIC, pelletized, and sent to the UC Davis Stable Isotope Facility for determination of carbon content and ^13^C isotopic abundance. Rates were integrated over the depth of the euphotic zone to determine integrated NPP (i.e., mg C m^-2^ d^-1^). Total integrated and size-fractionated chlorophyll-*a* (chlA) was measured as in Strom et al. (2016) and integrated to 75 m.

### Gel trap imaging and image processing

Polyacrylamide gels containing intact sinking particles were imaged in darkfield illumination with oblique lighting from multiple angles according to O’Daly et al. (O’Daly et al. 2020). A ruler positioned at the same height as the surface of the gel was imaged using the same settings for each gel to calculate pixels per centimeter for each imaged gel. Gel images were cropped to remove lighting artifacts and jar edges before being processed to detect particles according to Durkin et al. (Durkin et al. 2021) with a few modifications (Fig. 2, panel A). Image processing methods were modified to better select particles and minimize false-positive selections. Background determination followed the regional maxima method using the *skimage.morphology.reconstruction* function in Python to create a background image, which was subtracted from the original image. Edge detection was used before thresholding to identify and label in-focus particle edges. In the brightness threshold step, the triangle method was used to select the best thresholding value for each image, rather than a fixed thresholding value, by using the *skimage.filters.threshold_triangle* function. Binary processed images were saved as JPEGs (Fig. 2, panel B) and loaded into Adobe Photoshop CS6 for manual validation. Particles that were incorrectly segmented were manually corrected. The image was then saved and reloaded into the Python routine. A one-pixel erosion step was added before identifying and matching in-focus particle edges to remove a halo effect around particles. Finally, particles were counted and measured (Fig. 2, panel C). Calculating particle fluxes in units of # m^-2^ d^-1^ occurred by grouping particles by their equivalent spherical diameter (ESD) into logarithmically spaced size bins with edges at 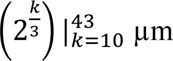 (Durkin et al. 2021). Uncertainty was calculated assuming a Poisson distribution in particle counts.

**Fig. 2.**
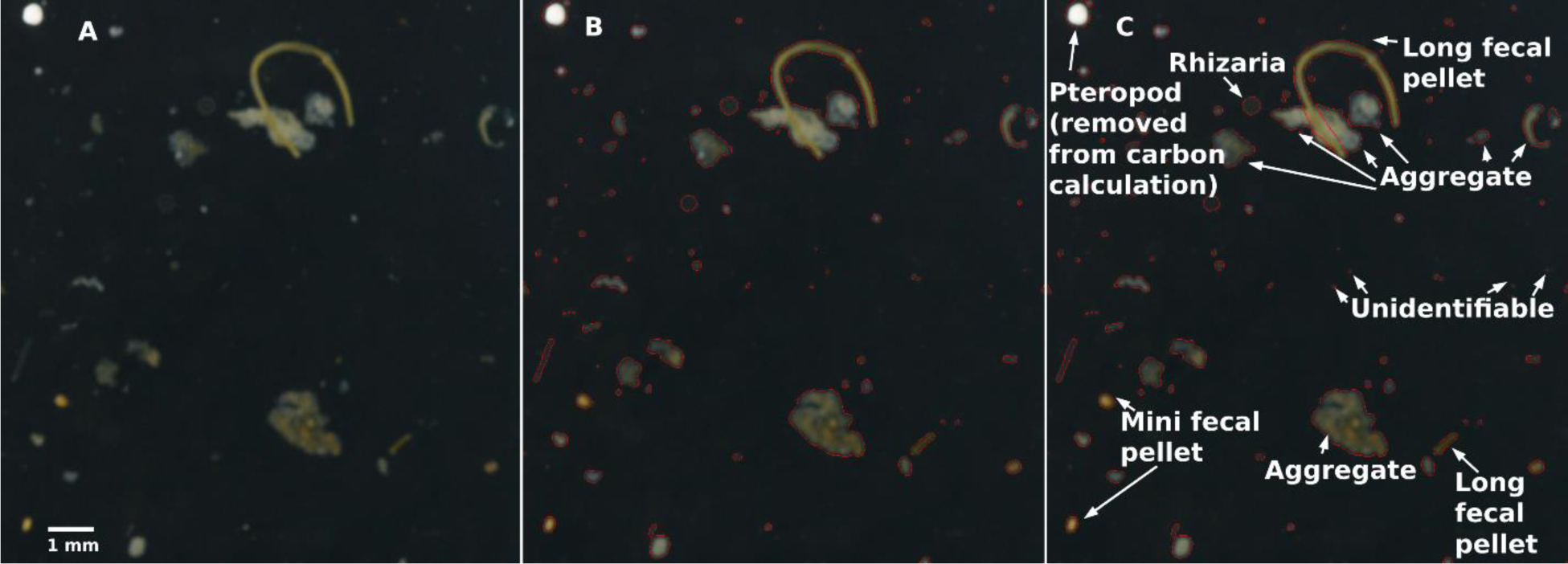
Particle segmentation and identification. (A) Sinking particle in polyacrylamide gel imaging, (B) automatic segmentation, (C) manual correction to segmentation, and classification.

### Gel trap carbon content by particle type

Particles were categorized into ten different particle types (Fig. 2, panel C): aggregates, dense detritus, long fecal pellet, large loose fecal pellet, short fecal pellet, salp fecal pellet, mini pellet, rhizaria, phytoplankton, and unidentifiable as in Durkin et al. (Durkin et al. 2021). Rhizaria were included in the carbon flux modeling because most rhizaria can be considered part of the gravitational flux (Bernstein et al. 1987; Michaels et al. 1995; Lampitt et al. 2009). All mesozooplankton were identified in images but removed from consideration when modeling carbon flux from the identified particles because they were assumed not to be part of the gravitational sinking flux. Particles manually identified from GAK15 were used as a training set in a model to predict the type of the remaining particles using high-level Keras preprocessing utilities from the Tensorflow package in Python to speed up the manual validation of all particles. Identification becomes more challenging as particle size decreases. More than 90% of particles smaller than 100 µm were unidentifiable while only ∼23% of particles larger than 100 µm were unidentifiable.

We calculated the carbon content for each particle type using a hybrid of inverse modeling and published values depending on particle type. Particle volume was calculated from ESD using type-specific ESD to volume relationships (Supporting Information Table 2) (Durkin et al. 2021). Then particle volumes were converted to carbon using Equation 2 where C is the calculated carbon content per particle (mg C), A is a scaling coefficient (mg C μm^-3^), V is the calculated particle volume (µm^3^), and B is a unitless exponent parameter.

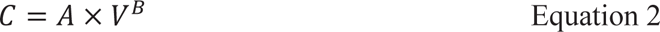

**Table 2.** Net primary productivity, total integrated chlorophyll-*a*, fraction of chlorophyll-*a* >20 µm, particulate organic carbon (POC) export flux, and export ratios at nine stations in the Northern Gulf of Alaska.

| Station | Euphotic Zone Depth (m) | Trap Depth (m) | Net Primary Productivity ($\text{mg C m}^{-2} \text{d}^{-1}$) | Total integrated chlorophyll-*a* ($\text{mg m}^{-2}$) | Fraction of chlorophyll-*a* > 20 $\mu\text{m}$ | POC Export Flux ($\text{mg C m}^{-2} \text{d}^{-1}$) | Export Ratio |
| --- | --- | --- | --- | --- | --- | --- | --- |
| GAK1 | 25 | 27 | 423 | 27.4 | 0.32 | $398 \pm 56$ | $0.94 \pm 0.13$ |
| GAK5 | 29 | 27 | 614 | 17.5 | 0.06 | $360 \pm 50$ | $0.59 \pm 0.08$ |
| GAK9 | 25 | 27 | 946 | 16.7 | 0.04 | $320 \pm 45$ | $0.34 \pm 0.05$ |
| GAK15 | 31 | 27 | 807 | 30.1 | 0.10 | $232 \pm 32$ | $0.29 \pm 0.04$ |
| KOD5 | 25 | 27 | 913 | 38.7 | 0.34 | $1039 \pm 145$ | $1.14 \pm 0.16$ |
| KOD10 | 37 | 53 | 698 | 8.9 | 0.08 | $482 \pm 67$ | $0.69 \pm 0.10$ |
| MID5 | 26 | 25 | 258 | 17.5 | 0.02 | $158 \pm 22$ | $0.61 \pm 0.09$ |
| MID10 | 34 | 25 | 526 | 25.2 | 0.32 | $346 \pm 48$ | $0.66 \pm 0.09$ |
| PWS2 | 22 | 80 | 240 | 58.3 | 0.17 | $172 \pm 24$ | $0.72 \pm 0.10$ |

Aggregates and unidentifiable particles were assumed to have the same A (A_agg_) and B (0.8) values (Menden-Deuer and Lessard 2000; Durkin et al. 2021). Dense detritus, large loose fecal pellets, long fecal pellets, short fecal pellets, and mini fecal pellets were assumed to have the same A value (A_FP_) and a B value of 1 (Wilson et al. 2008; Durkin et al. 2021). B values less than one are consistent with less efficient packaging of larger particles (Alldredge 1998). This pattern is more pronounced for aggregates and unidentifiable particles than fecal pellets since aggregate formation allows for large, loosely-packed particles (Johnson et al. 1996) while fecal pellets exhibit a more uniform carbon density (Bishop et al. 1980, 1986). Empirically derived or modeled A and B values were used for salp fecal pellets (Silver and Bruland 1981; Iversen et al. 2017), rhizaria (Menden-Deuer and Lessard 2000; Stukel et al. 2018), and phytoplankton (Menden-Deuer and Lessard 2000). Two parameters (i.e., A_agg_ and A_FP_) were fit to 26 observations. A maximum likelihood approach (i.e., Markov chain Monte Carlo) was used to fit the log-transformed carbon fluxes to the image-based analysis yielding both best-fit values and uncertainty (van Oevelen et al. 2010; Yingling et al. 2022). The *optimize.fmin* function in Python was used to determine starting values for the Markov chain Monte Carlo. A scale value of 1e-11 was used for A_agg_ and A_FP_. An exponent scale of 5 was used also. After burn-in, a final solution set containing 100,000 parameterizations was analyzed. The median of these values is considered the best-fit value. Uncertainty is reported as the 95% confidence interval around the median value (Supporting Information Fig. 2).

Twenty-six gel images were analyzed to identify particles and calculate gel-derived carbon flux estimates. Some gels, especially many at KOD5 and KOD10, were overloaded with particles and were excluded from the image processing step. Additionally, stations MID5 and PWS2 had POC flux that increased with depth, which may be a sign of non-steady state conditions: high lateral advection or resuspension from the seafloor. These stations were excluded from image processing. The gel-derived carbon flux estimates were within the expected uncertainty of the chemically derived carbon flux values, as most of the spread in the data is within the 2:1 and 1:2 lines (Supporting Information Table 3 and Supporting Information Fig. 3) (Durkin et al. 2021).

**Fig. 3.**
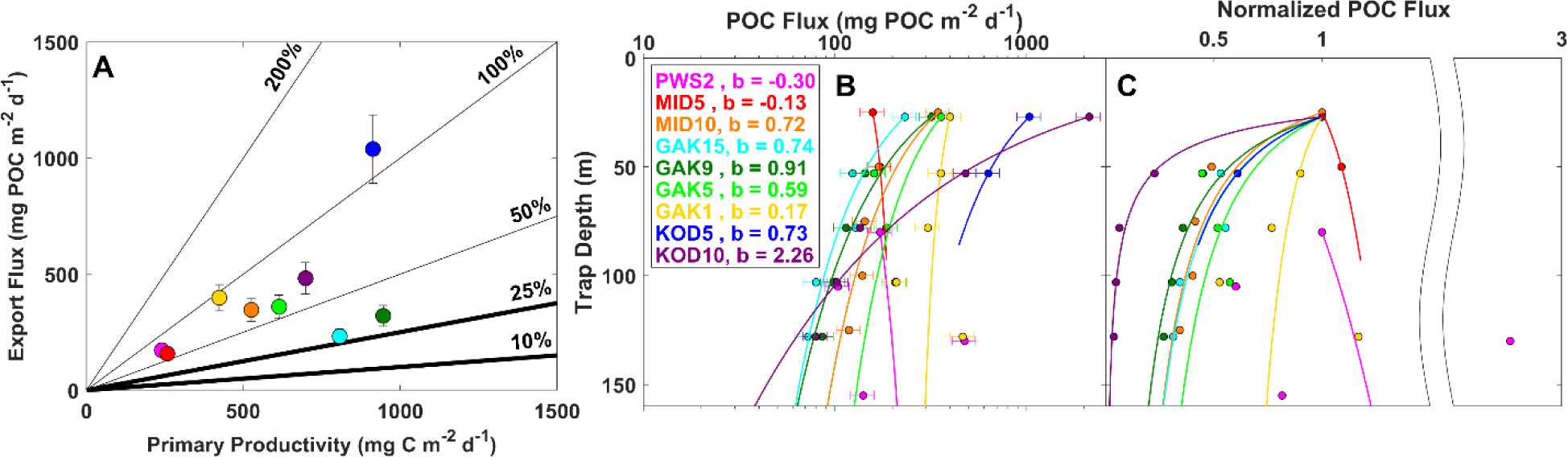
Strength and efficiency of the biological carbon pump. (A) Chemically-derived bulk **s**inking particulate organic carbon fluxes from drifting sediment traps and net primary productivity rates. Vertical error bars represent the average tube-to-tube standard deviation of about 14%. Diagonal lines show constant export ratios. Typical open ocean (10%) and typical shelf (25%) export ratio values are shown in bold (Dunne et al. 2007). (B) Chemically derived bulk sinking particulate organic carbon (POC) fluxes are shown over depth along with the best-fit attenuation curve and coefficient. Horizontal error bars represent the average tube-to-tube standard deviation of about 14%. (C) Normalized chemically-derived sinking POC fluxes from drifting sediment traps over depth.

## Particle concentration size distribution and sinking velocity

The UVP was used to calculate particle concentrations and size distributions throughout the water column. Particles from 102 µm to 26 mm were grouped into 5 m depth bins and one of 25 logarithmically spaced size bins with edges at 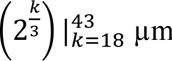. We calculated depth-specific particle abundance by dividing the number of particles per depth bin by the volume of water imaged per depth bin (#/L).

Size-specific average sinking velocity was calculated by dividing the flux of particles in each size bin by the concentration of particles in each size bin (Equation 3)

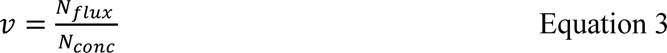

where v is the size-specific sinking velocity (m d^-1^), N_flux_ is the number fThelux of particles in a size bin (# particles in a size bin m^-2^ d^-1^) from the gels, and N_conc_ is the number concentration of particles in a size bin (# particles in a size bin m^-3^) from the UVP. This method averages the sinking velocity for all particles (sinking and suspended) in each size bin and can be thought of as apparent size-specific sinking velocity.

### Statistical analysis

A principal component analysis (PCA) was performed to compare stations by particle abundance and composition. The covariance matrix was calculated using the function *PCA* in the FactoMineR package in R. Quantitative supplementary variables calculated at each station were plotted in the same space. Variables included gel-estimated carbon flux (Est. C flux), chemically derived carbon flux (Chemical C flux), UVP-derived number concentration averaged from the 50 m above the trap (Total # concen), gel-derived number flux in (Total # flux), average UVP-and gel-derived sinking velocity averaged across sizes (Avg. sinking velocity; Equation 3), relative contribution of fecal pellet carbon flux (% FP flux), chemically-derived carbon to gel-derived volume ratio (C:V Ratio), slope of the gel-derived particle size distribution of sinking particles (Slope of PSD) calculated as the linear slope of the number flux versus size in each gel trap, depth of the trap (depth), and average sinking particle size (Mean size) calculated as the average particle size from each gel.

## Results and discussion

We studied the biological carbon pump (BCP) in the Northern Gulf of Alaska (NGA) during the summer of 2019 using drifting sediment traps to measure particulate organic carbon (POC) and particle type flux and 24-h deck-board ^13^C incubations to measure net primary productivity (NPP). This study sets a baseline for BCP in the NGA, as these are the first measurements of carbon export in this region of which we are aware.

### Net production of carbon in the euphotic zone

We measured NPP to determine the near-term input of carbon to the system. NPP was variable (Table 2), ranging from a low of 240 mg C m^-2^ d^-1^ in Prince William Sound (PWS2) to a high of 946 mg C m^-2^ d^-1^ near the shelf break (GAK9). NPP tended to increase with distance from shore, as on the GAK lines and MID lines (Table 2, Fig. 3, Fig. 4). This is expected, as higher rates of NPP are often observed at the shelf break where higher iron coastal waters meet higher nitrogen offshore waters. In addition, a shelf break front can form in the vicinity of the 300-m isobath (Fig. 1) promoting vertical transport of deeper nutrient-rich waters to the surface (Aguilar-Islas et al. 2016; Strom and Fredrickson 2020). The Kodiak Island line often behaves differently because of unique circulation patterns associated with the shallow banks on the shelf, which leads to less predictable spatial patterning. Observed NPP values and chlorophyll concentrations were typical for the NGA during summer (Strom and Fredrickson 2020), despite significantly higher sea surface temperatures and greater abundances of small nanoflagellates and picophytoplankton during the summer of 2019 (Cohen 2022). The fraction of chlA > 20 µm ranged from 0.04 at GAK9 to 0.34 at KOD5 and averaged 0.16 (Table 2).

**Fig. 4.**
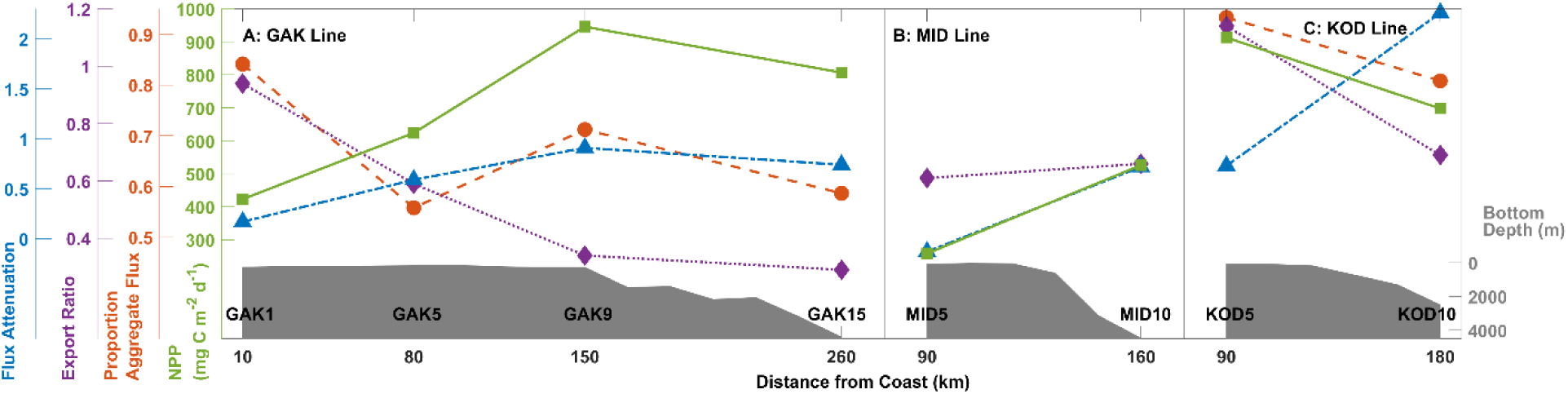
Offshore gradient in biological carbon pump (BCP) properties. Flux attenuation (blue triangles), export ratio (purple diamonds), proportion aggregate flux (orange circles), and net primary productivity (NPP; green squares) along with underlying bathymetry (filled grey patch) are plotted over distance from the coast for the (A) Gulf of Alaska (GAK) line, (B) Middle Island (MID) line, (C) Kodiak (KOD) line.

Overall, NPP estimates in the NGA averaged 603 ± 263 mg C m^-2^ d^-1^, ± 1 SD (Table 2, Fig. 3 panel A). For comparison, summer measurements in the offshore and high nutrient-low chlorophyll (HNLC) North Pacific at Ocean Station Papa (50 N 145 W) were similar to or less than the NGA: ∼160 mg C m^-2^ d^-1^ (Estapa et al. 2021), 700 mg C m^-2^ d^-1^ (Buesseler and Boyd 2009) and 300 – 1500 mg C m^-2^ d^-1^ (Welschmeyer et al. 1993). We would expect similar summertime NPP at Ocean Station Papa and the NGA because both systems are nutrient-limited at this time, the former by dissolved iron and the latter by macronutrients, and characterized by communities of pico-and nanophytoplankton (Booth et al. 1993; Welschmeyer et al. 1993; Strom et al. 2006). When compared to a more productive, coastal, upwelling region, the California Current Ecosystem (CCE) (32 – 35N, 120 – 124 W) has an average NPP similar to the NGA (747 ± 595 mg C m^-2^ d^-1^, ± 1 SD) yet with a maximum NPP more than twice that of the maximum measured in the NGA (2333 mg C m^-2^ d^-1^) (California Current Ecosystem LTER and Goericke 2022). See Supporting Information for a description of how the meta-analysis of the CCE data was analyzed.

### Strength of the biological carbon pump

To determine the strength of the BCP in the Northern Gulf of Alaska we deployed sediment traps and measured NPP at nine locations ranging over the continental shelf and slope during the summer of 2019 (Fig. 1, Table 1, Table 2). Overall, particulate organic carbon (POC) export flux was high, ranging from 158 to 1039 mg C m^-2^ d^-1^ and averaging 390 ± 265 mg C m^-2^ d^-1^ (± 1 SD) (Table 2). The magnitude of POC export flux varied considerably by station but was generally higher near the coast and diminished offshore (Table 2). POC export flux at the shelf stations ranged from 158 to 1039 mg C m^-2^ d^-1^ and averaged 408 ± 324 mg C m^-2^ d^-1^ (± 1 SD) (Table 2). Meanwhile, POC export flux at the slope stations ranged from 232 to 482 mg C m^-2^ d^-1^ and averaged 353 ± 125 mg C m^-2^ d^-1^ (± 1 SD). Along the GAK and KOD lines at all depths, there was a clear decrease in the magnitude of POC flux progressing offshore.

To put this in context, summertime POC export flux estimates at Ocean Station Papa were 9.5 - 78.2 mg C m^-2^ d^-1^ (Estapa et al. 2021) and 97 mg C m^-2^ d^-1^ (Buesseler and Boyd 2009), about one-fifth and one-third, respectively, of the average POC export flux in the NGA. In the CCE, POC fluxes ranged from 42 to 437 mg C m^-2^ d^-1^ and averaged 185±121 mg C m^-2^ d^-1^ (± 1 SD), with a median of 149 mg C m^-2^ d^-1^ (California Current Ecosystem LTER et al. 2022). Despite having similar or higher productivity, the average POC flux in the CCE was about half that of the NGA, and the maximum POC flux measured in the CCE was also about half of the highest POC export flux measured in the NGA. See Supporting Information for a description of CCE data meta-analysis methods. Overall, we measured a strong BCP during the summer in the NGA.

This study occurred exclusively during the summer of 2019 in the NGA: a system known for high seasonal variability (Waite and Mueter 2013) and high interannual variability (Strom et al. 2016) in chlorophyll and primary production. While we do not have previous carbon flux measurements from this region, high seasonal variability in carbon fluxes has been measured at Ocean Station Papa (Timothy et al. 2013). Therefore, it is likely that there is high seasonal and interannual variability in POC fluxes in the NGA.

PIC, PN, C:N, δ^13^C, δ^15^N, and total SPM fluxes are presented in the supporting information (Supporting Information Table 2, Supporting Information Fig. 4).

### Efficiency of the biological carbon pump

The export ratio (i.e., export ratio = export flux / NPP) quantifies the efficiency of the BCP, with higher export ratios indicating a more efficient BCP. Export ratios ranged from 29 - 114% (average 67%, median 65%) indicating that POC was efficiently transported out of the euphotic zone (Table 2, Fig. 3, panel A). Stations with a high export ratio and relatively low primary productivity (e.g., GAK1 in yellow) can potentially indicate advected material entering the water column, non-steady-state conditions, or a collection bias. Export ratios tended to exhibit an inverse relationship to NPP and flux attenuation, being higher on the shelf and decreasing offshore, and a positive relationship with export flux (Fig. 4). The MID line showed nearly identical export ratios at MID5 and MID10.

Overall, observed export ratios were very high (average 67%) compared to typical export ratios of around 25% near the shelf (Dunne et al. 2007; Henson et al. 2015; Kelly et al. 2018). Global models predict export ratios of 0.1 to 0.15 for the NGA (Henson et al. 2012), comparable to other measurements in the North Pacific. During summer, export ratios measured at Ocean Station Papa ranged from 6 – 18% and averaged 10% (Estapa et al. 2021) and were 14% in a previous study (Buesseler and Boyd 2009), about five times less than the average export ratio observed in the NGA. This pattern is unexpected, as Ocean Station Papa and the NGA share similar summertime nutrient limitation, rates of NPP, and phytoplankton community composition. However, the NGA is closer to the coast and coastal areas generally are more associated with higher carbon flux and export efficiency (Dunne et al. 2007). In the CCE, a more coastal region than the NGA, summer export ratios ranged from 10 – 42% and averaged 30±11% (± 1 SD), with a nearly identical median of 31% (see Supporting Information for how these values were calculated). The average export ratio in the CCE was about half of that in the NGA despite the CCE being a high-productivity, high-flux site. Our data suggest that the summertime BCP in the NGA is more efficient than previously assumed.

We expect that communities dominated by large cells will have higher export efficiency. Pico-and nanoplankton typically dominate the community during summertime, with 2019 as no exception (Cohen 2022). However, the proportion of chlA > 20 µm (microphytoplankton and larger) significantly positively correlated with export ratios (Pearson correlation coefficient, r(7) = .49, p = .02) (Fig. 5). The highest proportion of chlA > 20 µm observed was only around one third. This suggests that larger phytoplankton are driving carbon export efficiency, even though they only make up a minority percentage of total chlA. Pico and nanoplankton are thought to contribute to less efficient export due to their smaller size and lack of ballasting (Stokes 1851; Michaels and Silver 1988). However, other studies suggest that picoplankton can be exported efficiently if aggregates are formed (Richardson and Jackson 2007; Richardson 2019). Our data support the hypothesis that communities dominated by small cells can still have efficient export, but more efficient export is likely when the proportion of large cells is higher.

**Fig. 5.**
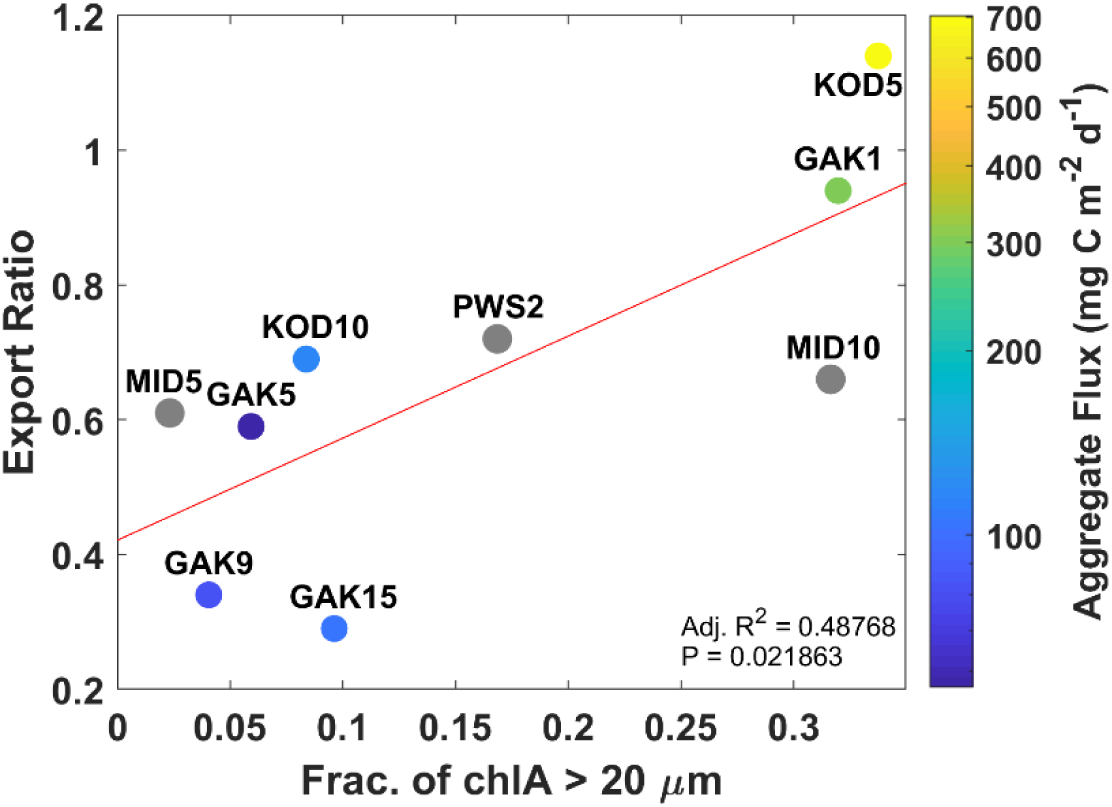
Drivers of efficient export. Fraction of chlorophyll-*a* (chlA) > 20 µm significantly correlates with export ratio (Pearson correlation coefficient, r(7) = .49, p = .02) colored by aggregate flux (in mg C m^-2^ d^-1^). Linear regression between the fraction of chlA > 20 µm and export ratio is shown as the red line.

POC fluxes generally decreased over depth consistent with a power-law curve (Fig. 3, panels B and C), with an exponent (*b*) describing how efficiently POC is transported from the base of the euphotic zone into the deep ocean (e.g., originally proposed as *b* = 0.86) (Martin et al. 1987; Buesseler et al. 2007). In our data, *b* ranged from -0.3 to 2.26, with lower values indicating a more efficient transfer of POC from the euphotic zone into the deep ocean. Our observations were lower (mean: 0.63 ± 0.74, ± 1 SD) than many typically observed (e.g., 0.86 in the northeast Pacific (Martin et al. 1987), 1.16 at Ocean Station Papa (Buesseler and Boyd 2009), 1.33 at station ALOHA (Buesseler et al. 2007)). It is somewhat unexpected that we measured such low POC flux attenuation in the NGA, as higher flux areas, i.e., coastal areas and higher latitude systems, often are modeled to have higher POC flux attenuation (Henson et al. 2012; Marsay et al. 2015). However, this topic is still being debated, as other modeling studies find more efficient BCP in high-latitude systems (Weber et al. 2016). Our observed, rather than modeled, POC flux attenuation measurements provide important evidence of an efficient BCP in a high-latitude coastal system.

POC flux attenuation also varied spatially in our study. PWS2 and MID5 both had negative attenuation coefficients, indicating an increase in carbon flux with depth. MID5 was the shallowest station and therefore may have been experiencing resuspension from the seafloor. The shallowest trap at PWS2 was 80m, far below the actual euphotic zone depth of 22 m. This substantially increases the uncertainty of the best-fit *b* value. Overall, carbon flux attenuation increased with distance from shore along all lines and had a similar pattern as NPP (Fig. 4).

### Contribution of carbon flux by particle types

To calculate the contribution of POC flux by particle type, we imaged the sinking particles, segmented those particles, identified particle types, and calculated particle volume by equivalent spherical diameter (ESD). Finally, we modeled A and B values using Equation 2 to find the best fit between the chemically derived POC flux and gel-estimated POC flux. Using this method, we can determine the contribution of each particle type (i.e., aggregates, fecal pellets, etc.) to the total POC flux.

The best-fit value A_agg_ for aggregates and unidentifiable particles was 1.482 x 10^-10^ mg C µm^-3^ with a 95% confidence interval of 0.684 – 2.189 x 10^-10^ mg C µm^-3^ (Supporting Information Table 2). The best-fit A_FP_ value for dense detritus, long, large loose, short, and mini fecal pellets was smaller than A_agg_ (1.196 x 10^-11^ mg C µm^-3^ with a 95% confidence interval of 0.059 – 3.826 x 10^-11^ mg C µm^-3^). This smaller A value is paired with a larger B value of 1 (as compared to 0.8), such that aggregates and fecal pellets of the same ESD have carbon contents of the same order of magnitude over the size range we analyze from the polyacrylamide gels (∼20 – 3000 µm). These values agree well with previous empirically-or model-derived values. See Supporting Information for a mini-review of these model results compared to empirical values (Supporting Information text, Supporting Information Fig. 5, Supporting Information Table 4).

The majority of the POC flux at nearly all stations and depths comes from aggregates, ranging from 41 - 93% and averaging 61 ± 13% (± 1 SD) (Fig. 6, Supporting Information Table 5). This majority further increases when considering all non-fecal pellet sources, the combined contribution of aggregates, dense detritus, and unidentifiable particles, ranging from 54 - 96% and averaging 74 ± 11% (± 1 SD). Carbon from fecal pellets made up a small percentage of sinking carbon flux, ranging from 4 - 45% and averaging 25 ± 10% (± 1 SD). Both overall and relative carbon flux from short fecal pellets, likely coming from small copepods (e.g., *Pseudocalanus*) or larvaceans (Wilson et al. 2008), increased with distance from shore. Long fecal pellets, likely from large copepods (e.g., *Neocalanus* spp.) or euphausiids (Wilson et al. 2008), were the most important fecal pellet type for carbon flux and relative carbon flux at all stations except KOD5 where large loose pellets, mini pellets, and short pellets all contributed higher carbon flux and relative flux than long fecal pellets. Ninety-five percent of the variability in overall gel-derived carbon flux was due to variability in aggregate carbon flux. The relative contribution of aggregates decreased moving offshore on the KOD line and had a similar pattern on the GAK line with some variability (Fig. 4). Aggregates contributed the most to carbon flux at GAK1, followed by GAK9, GAK15, and GAK5. Overall, we observed an efficient BCP with high export ratios and a predominance of aggregate flux.

**Fig. 6.**
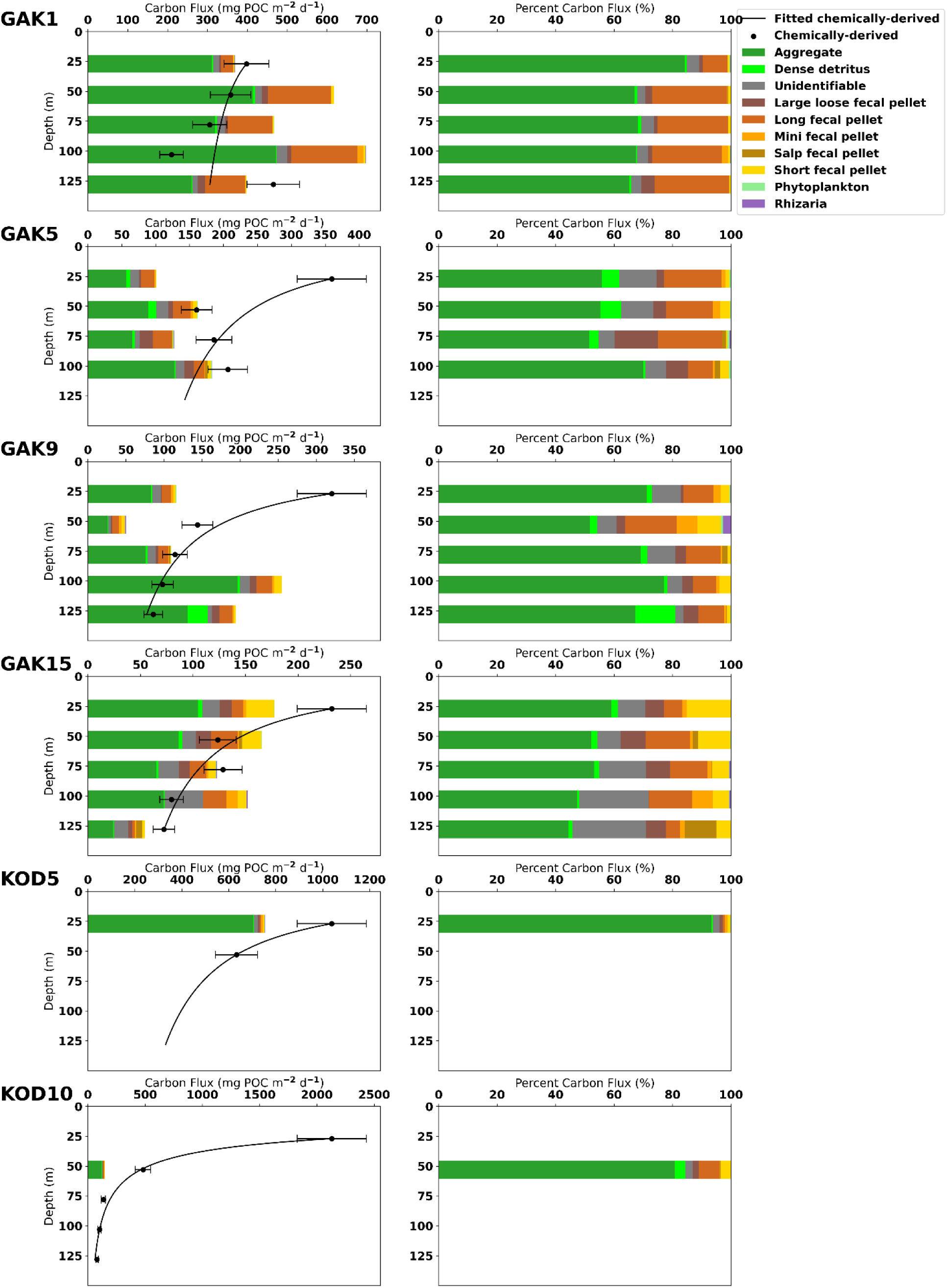
Contribution of carbon flux by particle type. Left columns: Depth profiles of gel-derived particulate organic carbon (POC) flux from June and July 2019 in the Northern Gulf of Alaska (NGA) by each particle type at the indicated locations. Black circles show the corresponding chemically-derived bulk POC fluxes with POC attenuation shown in black. Horizontal error bars indicate an average tube-to-tube error of about 14%. Right columns: Depth profiles of relative POC flux by particle type from gel analysis.

Most particles identified as aggregates and dense detritus appeared to be made up of largely green phytoplankton-derived material. Almost no particles were identified as larvacean houses, which have a much lower carbon-to-volume ratio than most aggregates, even though larvaceans were more abundant than normal in the Northern Gulf of Alaska during the summer of 2019 (R. Hopcroft and E. Stidham, pers. comm.).

GAK1, GAK5, and GAK9 showed some signs of increasing gel-derived POC flux over depth, while the chemically-derived POC fluxes decreased with depth at these stations (Fig. 6). This could be explained by carbon-to-volume ratios that change with depth. As particles sink and undergo remineralization, their mass may decrease while their ESD remains the same. Future studies should explore this idea to determine whether improving the fit between chemically-and gel-derived POC fluxes is possible.

### Drivers of carbon export

To determine if certain particle types were correlated with higher carbon export, a PCA was performed on the carbon flux by particle type at each station/depth sample (Fig. 7, panel A). Aggregate carbon flux is strongly associated with total carbon flux, phytoplankton flux, and long fecal pellet flux (Fig. 7, panel B). Gel-estimated and chemically derived carbon flux tended to decrease as stations moved offshore (Fig. 7, panel C). Total number concentration (i.e., ambient particle concentrations) and total number flux are both positively correlated to carbon flux. Interestingly, sinking velocity, mean sinking particle size, and trap depth are poor indicators of carbon flux. The carbon-to-volume ratio has an inverse relationship with carbon flux. Estimated carbon flux from aggregates, long fecal pellets, and phytoplankton are significantly correlated (linear regression, F-statistic, p < 0.05) with the total carbon flux, however, carbon flux from aggregates most strongly correlates with carbon flux.

**Fig. 7.**
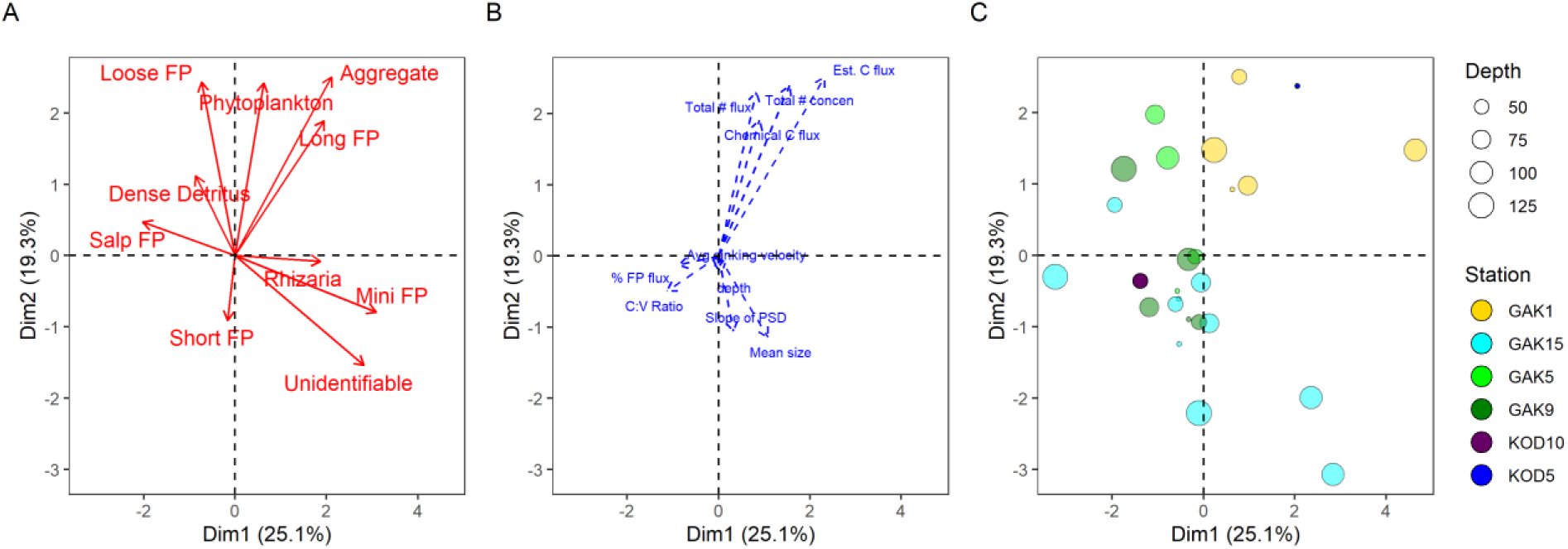
(A) PCA on the carbon flux by particle type at each station/depth location, shown in red arrows: FP = Fecal Pellet. (B) Quantitative supplementary variables estimated at each depth/station pair: total gel-estimated carbon flux (Est. C flux), total chemically-derived particulate organic carbon flux (Chemical C flux), total number concentration averaged from the 50 m above the trap depth (Total # concen), gel-derived total number flux (Total # flux), average sinking velocity (Avg. sinking velocity), relative contribution of fecal pellets to carbon flux (% FP flux), calculated carbon to volume ratio (C:V Ratio), slope of the gel-derived particle size distribution of sinking particles (Slope of PSD), depth of the trap (depth), and average sinking particle size (Mean size). (C) Individuals are plotted as colored dots corresponding to a sample (i.e., one depth/station pair). Colors and size correspond with the station and depth at which the sediment trap sampled, respectively.

Efficient export in the NGA during summer is driven by a few parameters. Export ratio correlated significantly with the proportion of chlA > 20 µm, total aggregate flux, export flux, and percent aggregate flux (Pearson, p ≤ .05) (Fig. 5). Aggregate flux also significantly correlated with the proportion of chlA > 20 µm (Pearson, p ≤ .05) (Fig. 5). Interestingly, NPP does not significantly correlate with any other variable. KOD5 and GAK1, for example, had the highest export ratios, proportion of chlA > 20 µm, and contributions of aggregate flux. This provides a framework for conditions that allow for efficient export, besides efficient export driven by high fecal pellet flux as previously described (Ebersbach and Trull 2008; Laurenceau-Cornec et al. 2015).

To determine if certain particle types more efficiently transport carbon to depth, we performed linear regressions on carbon flux over depth from different stations and different particle types (Supporting Information Fig. 6). Certain particle types are not more likely to reach different depths than others. Only four of these carbon fluxes by station and particle type combinations have significant relationships over depth; at GAK15, aggregates decreased over depth with a slope of -0.67 (p = 0.025), dense detritus decreased over depth with a slope of -0.04 (p = 0.005), and short fecal pellets decreased over depth with a slope of -0.23 (p = 0.016); at GAK9 large loose fecal pellets increased over depth with a slope of 0.10 (p = 0.011). Within each particle type, the slopes also have mixed positive and negative relationships depending on the station. Therefore, these data do not strongly support a claim that carbon from certain particle types is more or less likely to reach deeper depths than other particle types.

Debate continues over whether zooplankton fecal pellet production through the consumption of phytoplankton and aggregates results in higher or lower carbon export (Turner 2015). In some studies, less dense, sometimes smaller material that is repackaged into larger, denser particles results in higher carbon export (Iversen and Lampitt 2020). In other studies, mineral ballasted aggregates exported carbon more effectively than fecal pellets (Armstrong et al. 2002), as fecal pellets can have high rates of particle-associated microbial respiration (Smetacek 1985; Ploug et al. 2008a; Kobari et al. 2013). Our results suggest that particle type did not strongly impact the efficiency at which POC was transferred from the euphotic zone to the deep ocean during summer in the NGA. One caveat is that our deepest trap was around 125 m, relatively shallow. It is possible that the particle type-specific effects only apply to the deeper ocean.

Performing a similar analysis against particle sizes, we found no significant difference in ESDs with depth, station, or type (Supporting Information Fig. 7). The absence of relationships between particle size and depth is consistent with three different explanations. First, if particle formation below the euphotic zone is minor compared to flux from the euphotic zone, then consistency of particle sizes would imply similar attenuation length scales across particle classes. Secondly, if particle formation below the euphotic zone is significant relative to euphotic zone export, then it implies that the size spectra formed in the twilight zone are similar to those in the euphotic zone, and perhaps that similar repackaging processes are dominant. Thirdly, it is possible that the particle size-specific effects are relatively slow compared to the residence time of sinking particles above our traps and that deeper traps may resolve such patterns.

### Marine Heatwave

During 2019 sea surface temperatures in the NGA were about 2°C warmer than the long-term mean (Danielson et al. 2022), characteristic of a marine heatwave. The 2019 marine heatwave was associated with higher abundances of picophytoplankton observed during the summer in the NGA (Cohen 2022). Generally, warmer ocean temperatures are thought to increase microbial respiration rates and decrease the sinking speed of particles generated by the food web (Michaels and Silver 1988; Vaqué et al. 2019). However, we measured a strong and efficient BCP characterized by a high proportion of aggregate flux during the summer in the NGA. If our observations of a strong and efficient BCP are representative of this system, this might imply that BCP in the NGA is even stronger and more efficient in non-heatwave summers.

## Conclusions

This study is the first description of the biological carbon pump (BCP) in the NGA. The BCP was strong and efficient during the summer of 2019. Net primary productivity (NPP) was typical of other past measurements in the subarctic and coastal North Pacific; however, POC fluxes and export ratios were much higher with lower carbon flux attenuation and higher contributions of aggregates, than prior results from similar regions (Buesseler and Boyd 2009; Durkin et al. 2021; Estapa et al. 2021). By using a comprehensive approach that brings together sediment trap sampling and imaging, optically measured distribution of sinking and suspended particles, and incubations, our data suggest that the main driver of carbon flux in the NGA during summer was aggregation processes and the main drivers of efficient carbon export were the proportion of chlorophyll-*a* in the large size fraction (>20 µm) and aggregation processes. These results lead us to question our expectations about what conditions and processes can create strong and efficient flux events noting that high and efficient flux events can occur with moderate contribution from zooplankton fecal pellets and communities dominated by pico-and nanophytoplankton. These results can help improve this region’s climate and ecological models to better predict the fate of organic material produced through photosynthesis.

## Acknowledgments

We thank the Captain and crew of the R/V *Sikuliaq*, the science party on the 2019 Summer NGA-LTER Process cruise, and Catherine Fuller, NOAA Teacher at Sea, for assistance with sample collection. We also thank Colleen Durkin for discussions of polyacrylamide gel image analyses. Funding for this effort was provided by NSF NGA-LTER grant number 1656070 and NSF CAREER Award grant number 1654663. The authors declare no conflicts of interest relevant to this study.

## Data Availability Statement

All data associated with this study are publicly available in the supporting information and on GitHub (https://github.com/shodaly2/ODaly_et_al_2023_L_and_O_NGA_Flux).

