## Supporting Information for "Strong and efficient biological carbon pump in the Northern Gulf of Alaska during summer"

***Title***

***Authors and affiliations***

Stephanie H. O'Daly[1*] (0000-0002-5031-6842),

Gwenn M. M. Hennon[1] (0000-0002-5232-3843),

Thomas B. Kelly[1] (0000-0001-6285-1925),

Suzanne L. Strom[2] (0000-0001-5878-4790),

Andrew M. P. McDonnell[1] (0000-0003-1408-4869),

[1]College of Fisheries and Ocean Sciences, University of Alaska Fairbanks, Fairbanks, Alaska

[2]Shannon Point Marine Center, Western Washington University, Anacortes, Washington

***Running head***

Strong and efficient summer BCP in NGA

***Keywords***

Export, carbon flux, POC flux, LTER, primary productivity, sediment trap, UVP, fecal pellets, aggregates, North Pacific

**Supporting Information for: O'Daly et al., 2023 Limnology and Oceanography**

**California Current Ecosystem biological carbon pump meta-analysis**

The California Current Ecosystem (CCE) (32 – 35N, 120 – 124 W) is a coastal site with frequent upwelling ~2500 km south of the NGA study region that is also a part of the Long Term Ecological Research program (Kahru et al. 2015). We found and analyzed 16 POC export flux-NPP rate measurement pairs from 2011 – 2014 from May to August (California Current Ecosystem LTER and Goericke 2022; California Current Ecosystem LTER et al. 2022). We averaged NPP rates when there was more than one reported during the time of the sediment trap deployment. If multiple inline sediment traps collected POC flux we selected the shallowest depth assuming this is the export depth.

**Model results compared to empirical values**

A mini-review of empirically derived and modeled A and B values reveals a large spread in A and B values for different particle types (Supporting Information Fig. 5, Supporting Information Table 4). Fecal pellets tend to have B values close to 1, while aggregates and phytoplankton tend to have B values smaller than fecal pellets' B values. Results from this study (shown in diamond markers) are within the range of previous empirically derived and modeled B values, however, A values from this study are slightly larger than previous empirically derived and modeled A values (Supporting Information Fig. 5, panel A). All A values where B equals 1 are plotted sequentially by type for clarity (Supporting Information Fig. 5, panel B). While there is some spread in the gel-derived carbon values, they match within known tube-to-tube variability in carbon flux and are close in magnitude to previously empirically derived and modeled values.

### Supporting Information Tables

Supporting Information Table 1. Formalin-preserved bulk particulate organic carbon (POC) at each cast, depth, and tube, with average at each cast and depth, residual, relative residual, absolute value of relative residual

| Station Number | Depth (m) | Observation (mg C) | Mean | Residual | Relative Residual | Absolute Value Relative Residual |
|---|---|---|---|---|---|---|
| PWS2 | 155 | 217.426974 | 241.5242 | -24.0972 | -0.09977 | 0.099771 |
| PWS2 | 155 | 265.6213942 | 241.5242 | 24.09721 | 0.099771 | 0.099771 |
| PWS2 | 130 | 1448.533022 | 817.7262 | 630.8068 | 0.771416 | 0.771416 |
| PWS2 | 130 | 186.919386 | 817.7262 | -630.807 | -0.77142 | 0.771416 |
| PWS2 | 105 | 189.8236361 | 178.2477 | 11.5759 | 0.064943 | 0.064943 |
| PWS2 | 105 | 166.6718357 | 178.2477 | -11.5759 | -0.06494 | 0.064943 |
| PWS2 | 80 | 290.9671696 | 296.0953 | -5.12814 | -0.01732 | 0.017319 |
| PWS2 | 80 | 301.2234451 | 296.0953 | 5.128138 | 0.017319 | 0.017319 |
| MID5 | 50 | 429.5593757 | 427.5769 | 1.982435 | 0.004636 | 0.004636 |
| MID5 | 50 | 425.5945051 | 427.5769 | -1.98244 | -0.00464 | 0.004636 |
| MID5 | 25 | 526.289151 | 392.0836 | 134.2056 | 0.342288 | 0.342288 |
| MID5 | 25 | 257.8779922 | 392.0836 | -134.206 | -0.34229 | 0.342288 |
| MID10 | 125 | 174.6219065 | 157.7614 | 16.86055 | 0.106874 | 0.106874 |
| MID10 | 125 | 140.9008 | 157.7614 | -16.8606 | -0.10687 | 0.106874 |
| MID10 | 100 | 219.6721134 | 185.303 | 34.36916 | 0.185476 | 0.185476 |
| MID10 | 100 | 150.9337942 | 185.303 | -34.3692 | -0.18548 | 0.185476 |
| MID10 | 75 | 214.4925578 | 190.674 | 23.81859 | 0.124918 | 0.124918 |
| MID10 | 75 | 166.8553708 | 190.674 | -23.8186 | -0.12492 | 0.124918 |
| MID10 | 50 | 270.1523213 | 225.7132 | 44.43908 | 0.196883 | 0.196883 |
| MID10 | 50 | 181.2741696 | 225.7132 | -44.4391 | -0.19688 | 0.196883 |
| MID10 | 25 | 427.4661791 | 460.8097 | -33.3435 | -0.07236 | 0.072359 |
| MID10 | 25 | 494.1532443 | 460.8097 | 33.34353 | 0.072359 | 0.072359 |
| GAK15 | 128 | 213.941163 | 225.5967 | -11.6555 | -0.05167 | 0.051665 |
| GAK15 | 128 | 237.252256 | 225.5967 | 11.65555 | 0.051665 | 0.051665 |
| GAK15 | 103 | 231.5189815 | 248.4882 | -16.9692 | -0.06829 | 0.06829 |
| GAK15 | 103 | 265.457396 | 248.4882 | 16.96921 | 0.06829 | 0.06829 |
| GAK15 | 78 | 567.418761 | 401.2393 | 166.1795 | 0.414165 | 0.414165 |
| GAK15 | 78 | 235.0598374 | 401.2393 | -166.179 | -0.41417 | 0.414165 |
| GAK15 | 53 | 329.4058668 | 385.5174 | -56.1115 | -0.14555 | 0.145549 |
| GAK15 | 53 | 441.6289647 | 385.5174 | 56.11155 | 0.145549 | 0.145549 |
| GAK15 | 27 | 554.8840967 | 724.0718 | -169.188 | -0.23366 | 0.233661 |
| GAK15 | 27 | 893.2594951 | 724.0718 | 169.1877 | 0.233661 | 0.233661 |

| | | | | | | |
|---|---|---|---|---|---|---|
| GAK9 | 128 | 228.9217612 | 247.1797 | -18.258 | -0.07387 | 0.073865 |
| GAK9 | 128 | 265.4376907 | 247.1797 | 18.25796 | 0.073865 | 0.073865 |
| GAK9 | 103 | 261.2150029 | 282.1576 | -20.9426 | -0.07422 | 0.074223 |
| GAK9 | 103 | 303.1002671 | 282.1576 | 20.94263 | 0.074223 | 0.074223 |
| GAK9 | 78 | 351.4848236 | 329.4212 | 22.06363 | 0.066977 | 0.066977 |
| GAK9 | 78 | 307.357565 | 329.4212 | -22.0636 | -0.06698 | 0.066977 |
| GAK9 | 53 | 485.4884127 | 414.1452 | 71.34326 | 0.172266 | 0.172266 |
| GAK9 | 53 | 342.8018982 | 414.1452 | -71.3433 | -0.17227 | 0.172266 |
| GAK9 | 27 | 672.7639203 | 921.7546 | -248.991 | -0.27013 | 0.270127 |
| GAK9 | 27 | 1170.745336 | 921.7546 | 248.9907 | 0.270127 | 0.270127 |
| GAK5 | 103 | 255.8325672 | 298.2629 | -42.4303 | -0.14226 | 0.142258 |
| GAK5 | 103 | 340.693214 | 298.2629 | 42.43032 | 0.142258 | 0.142258 |
| GAK5 | 78 | 335.6610049 | 268.5543 | 67.10675 | 0.249882 | 0.249882 |
| GAK5 | 78 | 201.4474982 | 268.5543 | -67.1068 | -0.24988 | 0.249882 |
| GAK5 | 53 | 213.8624964 | 231.6297 | -17.7672 | -0.07671 | 0.076705 |
| GAK5 | 53 | 249.3969136 | 231.6297 | 17.76721 | 0.076705 | 0.076705 |
| GAK5 | 27 | 557.1140395 | 519.5162 | 37.59788 | 0.072371 | 0.072371 |
| GAK5 | 27 | 481.9182812 | 519.5162 | -37.5979 | -0.07237 | 0.072371 |
| GAK1 | 128 | 677.4726863 | 618.3105 | 59.1622 | 0.095684 | 0.095684 |
| GAK1 | 128 | 559.1482768 | 618.3105 | -59.1622 | -0.09568 | 0.095684 |
| GAK1 | 103 | 271.5133261 | 278.7164 | -7.20303 | -0.02584 | 0.025844 |
| GAK1 | 103 | 285.9193854 | 278.7164 | 7.20303 | 0.025844 | 0.025844 |
| GAK1 | 78 | 372.4082064 | 406.3768 | -33.9686 | -0.08359 | 0.083589 |
| GAK1 | 78 | 440.345308 | 406.3768 | 33.96855 | 0.083589 | 0.083589 |
| GAK1 | 53 | 552.888948 | 476.0819 | 76.80705 | 0.161332 | 0.161332 |
| GAK1 | 53 | 399.2748449 | 476.0819 | -76.8071 | -0.16133 | 0.161332 |
| GAK1 | 27 | 439.0904847 | 529.0775 | -89.987 | -0.17008 | 0.170083 |
| GAK1 | 27 | 619.064471 | 529.0775 | 89.98699 | 0.170083 | 0.170083 |
| KOD5 | 53 | 663.895395 | 681.7883 | -17.8929 | -0.02624 | 0.026244 |
| KOD5 | 53 | 699.6811072 | 681.7883 | 17.89286 | 0.026244 | 0.026244 |
| KOD5 | 27 | 1054.851497 | 1117.892 | -63.0405 | -0.05639 | 0.056392 |
| KOD5 | 27 | 1180.932544 | 1117.892 | 63.04052 | 0.056392 | 0.056392 |
| KOD10 | 128 | 203.4807726 | 209.3364 | -5.85559 | -0.02797 | 0.027972 |
| KOD10 | 128 | 215.1919585 | 209.3364 | 5.855593 | 0.027972 | 0.027972 |
| KOD10 | 103 | 220.8036851 | 270.0172 | -49.2136 | -0.18226 | 0.182261 |
| KOD10 | 103 | 319.2307865 | 270.0172 | 49.21355 | 0.182261 | 0.182261 |
| KOD10 | 78 | 367.0459523 | 357.0135 | 10.03248 | 0.028101 | 0.028101 |
| KOD10 | 78 | 346.980989 | 357.0135 | -10.0325 | -0.0281 | 0.028101 |
| KOD10 | 53 | 1013.062828 | 1271.408 | -258.345 | -0.2032 | 0.203196 |
| KOD10 | 53 | 1529.753301 | 1271.408 | 258.3452 | 0.203196 | 0.203196 |
| KOD10 | 27 | 5200.770713 | 5620.063 | -419.293 | -0.07461 | 0.074606 |
| KOD10 | 27 | 6039.356176 | 5620.063 | 419.2927 | 0.074606 | 0.074606 |
| **Average** | | | | | | **0.141465** |

Supporting Information Table 2. Assumptions and equations used to model each particle type's gel-derived particulate organic carbon content. Format taken from Durkin et al. (2021).

| Classification | Shape | Width ($w$) | Length ($l$) | Volume ($v$) | $C = A \times V^B$ (Equation 2) | | |
|---|---|---|---|---|---|---|---|
| | | | | | A | B | Ref |
| Aggregate | Sphere | $w = ESD$ | $l = ESD$ | $V = \frac{4}{3} \times \pi \times \left(\frac{ESD}{2}\right)^3$ | $1.482 \times 10^{-10}$ | 0.8 | This study |
| Dense detritus | Sphere | $w = ESD$ | $l = ESD$ | $V = \frac{4}{3} \times \pi \times \left(\frac{ESD}{2}\right)^3$ | | | |
| Large loose fecal pellet | Cylinder | $w = \frac{553 \times ESD}{ESD + 996}$ | $l = \frac{\pi \times (\frac{ESD}{2})^3}{w}$ | $V = l \times \pi \times \left(\frac{w}{2}\right)^2$ | | | |
| Long fecal pellet | Cylinder | $w = \frac{187 \times ESD}{ESD + 424}$ | $l = \frac{\pi \times (\frac{ESD}{2})^3}{w}$ | $V = l \times \pi \times \left(\frac{w}{2}\right)^2$ | | | |
| Short fecal pellet | Ellipsoid | $w = 0.54 \times ESD$ | $l = ESD$ | $V = \frac{4}{3} \times \frac{l}{2} \times \pi \times \left(\frac{w}{2}\right)^2$ | | | |
| Mini fecal pellet | Sphere | $w = ESD$ | $l = ESD$ | $V = \frac{4}{3} \times \pi \times \left(\frac{ESD}{2}\right)^3$ | $1.196 \times 10^{-11}$ | 1 | This study |
| Salp fecal pellet | Cuboid | $w = 0.63 \times ESD$ | $l = \frac{\pi \times (\frac{ESD}{2})^3}{w}$ | $V = l \times \pi \times \left(\frac{w}{4}\right)$ | $0.04 \times 10^{-9}$ | 1 | 1,2 |
| Rhizaria | Sphere | $w = ESD$ | $l = ESD$ | $V = \frac{4}{3} \times \pi \times \left(\frac{ESD}{2}\right)^3$ | $0.004 \times 10^{-9}$ | 0.939 | 3,4 |
| Phytoplankton | Cylinder | $w = \frac{187 \times ESD}{ESD + 424}$ | $l = \frac{\pi \times (\frac{ESD}{2})^3}{w}$ | $V = l \times \pi \times \left(\frac{w}{2}\right)^2$ | $0.228 \times 10^{-9}$ | 0.811 | 3 |
| Unidentifiable | Sphere | $w = ESD$ | l = ESD | $V = \frac{4}{3} \times \pi \times \left(\frac{ESD}{2}\right)^3$ | $1.482 \times 10^{-10}$ | 0.8 | This study |

Note. Parameters for Equation 2 were used from various reference studies: 1: Silver and Bruland (1981), 2: Iversen et al. (2017), 3: Menden-Deuer and Lessard (2000), 4: Stukel et al. (2018). ESD = equivalent spherical diameter.

Supporting Information Table 3. Formalin-preserved bulk particulate organic carbon (POC) flux, particulate inorganic carbon (PIC) flux, particulate nitrogen (PN), delta 13C, delta 15N, C:N, and total suspended particulate matter (SPM), average, and standard deviation (STD) at each cast and depth (see Table 1 for the cast to station translation)

| **Station** | **Depth (m)** | **Tube 1 POC Flux (mg C $m^{-2}$ $d^{-1}$)** | **Tube 2 POC Flux (mg C $m^{-2}$ $d^{-1}$)** | **Average POC Flux (mg C $m^{-2}$ $d^{-1}$)** | **STD error average POC Flux (mg C $m^{-2}$ $d^{-1}$)** | **Average PIC Flux (mg C $m^{-2}$ $d^{-1}$)** | **STD error average PIC Flux (mg C $m^{-2}$ $d^{-1}$)** | **Average PN Flux (mg C $m^{-2}$ $d^{-1}$)** | **STD error average PN Flux (mg C $m^{-2}$ $d^{-1}$)** | **Average Delta 13C flux (0/00)** | **STD error average Delta 13C flux (0/00)** | **Average Delta 15N** | **STD error Average Delta 15N** | **Average C:N Ratio** | **STD Error C:N Ratio** | **Average Total Dried Weight (mg SPM $m^{-2}$ $d^{-1}$)** | **STD error Average Total Dried Weight (mg SPM $m^{-2}$ $d^{-1}$)** |
|---|---|---|---|---|---|---|---|---|---|---|---|---|---|---|---|---|---|
| PWS2 | 155 | 126.520338 | 154.564579 | 140.542458 | 19.8302726 | 189.641626 | 124.043913 | 16.9925645 | 5.99743071 | -24.0595 | 0.5635641 | 8.8635 | 1.14056324 | 8.27082094 | 0.38010318 | 4082.98484 | 150.870161 |
| PWS2 | 130 | 108.768031 | 842.89858 | 475.833306 | 519.108689 | 10.505595 | -159.275138 | 94.012704 | 113.756096 | -23.4245 | 1.94666497 | 3.192 | 9.64352228 | 5.0613724 | 1.62919703 | 5634.71305 | 1330.40051 |
| PWS2 | 105 | 96.9860207 | 110.458009 | 103.722015 | 9.52613416 | 202.292303 | 176.549712 | 12.6642034 | 1.85122995 | -24.5995 | 0.1873833 | -2.7885 | 12.8318668 | 8.19017284 | 0.17263597 | 7293.12257 | 1728.14912 |
| PWS2 | 80 | 169.313236 | 175.281344 | 172.29729 | 4.22008893 | 135.368443 | 16.4442565 | 22.3807217 | 0.63108483 | -24.3345 | 0.08414571 | 8.3595 | 1.2720851 | 7.69846892 | 0.03734997 | 6158.42132 | 1248.1077 |
| MID5 | 50 | 171.010299 | 172.603444 | 171.806871 | 1.12652355 | -11.0528272 | 49.2428641 | 33.7107751 | 1.68589187 | -24.4625 | 0.1873833 | 7.9395 | 0.69225754 | 5.09649722 | 0.05043849 | 5772.74361 | 984.970765 |
| MID5 | 25 | 103.619271 | 211.470928 | 157.5451 | 76.2626378 | 39.8436575 | -72.3521334 | 33.4328462 | 19.0379763 | -24.58 | 0.2503158 | 7.3865 | 0.62437529 | 4.712285 | 0.74738446 | 4065.02943 | 1752.11146 |
| MID10 | 125 | 105.803533 | 131.12498 | 118.464257 | 17.9049672 | 26.5447081 | 37.9055133 | 16.8974386 | 2.46098789 | -24.5795 | 0.31607673 | 14.2685 | 8.63872355 | 7.01078189 | 0.20989473 | 7146.1408 | 2530.96708 |
| MID10 | 100 | 113.337388 | 164.95354 | 139.145464 | 36.4981306 | 50.9034347 | 14.7398189 | 20.6964701 | 8.49446268 | -24.219 | 0.30971277 | 6.896 | 1.72958319 | 6.72314957 | 0.48708883 | 6720.62628 | 725.661889 |
| MID10 | 75 | 125.293027 | 161.06417 | 143.178598 | 25.2940179 | -4.34878421 | -1.19132847 | 27.1325773 | 7.44797067 | -23.0075 | 0.08273149 | 9.19 | 2.84115505 | 5.27699955 | 0.32643652 | 8447.7146 | 3380.52246 |
| MID10 | 50 | 136.120218 | 202.85953 | 169.489874 | 47.1918201 | 117.797138 | 4.50040864 | 25.4316971 | 10.0823371 | -23.97 | 0.65760931 | 11.0705 | 2.95358503 | 6.6645129 | 0.48445485 | 7859.50336 | 3575.21223 |
| MID10 | 25 | 320.987758 | 371.063607 | 346.025683 | 35.408972 | 92.247515 | -1.73589992 | 62.8410583 | 5.63895111 | -24.0115 | 0.33870415 | 8.9155 | 1.39229325 | 5.50636307 | 0.13610156 | 8109.80602 | 707.962818 |
| GAK15 | 128 | 68.6016139 | 76.0764662 | 72.33904 | 5.28551876 | 5.96386141 | 2.75516969 | 9.66347244 | 0.99328371 | -25.0825 | 0.06717514 | 6.311 | 0.38466609 | 7.48582256 | 0.12611062 | 1630.00362 | 22.673835 |
| GAK15 | 103 | 74.2380547 | 85.1206263 | 79.6793405 | 7.69514011 | 24.4788452 | -2.0623863 | 9.96635303 | 1.39480053 | -25.62 | 0.10323759 | 6.5365 | 0.99489924 | 7.99483425 | 0.17003899 | 2217.87378 | 355.223415 |
| GAK15 | 78 | 75.373453 | 181.946486 | 128.659969 | 75.358514 | -31.1259323 | -68.8376634 | 18.1328461 | 9.51123891 | -25.381 | 0.14283557 | 6.2285 | 0.07990307 | 7.09540956 | 0.78625618 | 2485.08749 | 385.455195 |
| GAK15 | 53 | 105.626116 | 141.611176 | 123.618646 | 25.4452801 | 33.1614903 | 19.7059084 | 20.4508987 | 5.36543029 | -25.6125 | 0.15344217 | 8.575 | 2.94722106 | 6.04465591 | 0.33346645 | 2597.31725 | 241.85424 |
| GAK15 | 27 | 177.927165 | 286.429419 | 232.178292 | 76.7226796 | -49.4029844 | -44.2997462 | 45.5325085 | 15.807345 | -25.4545 | 0.21566757 | 5.0985 | 1.14763431 | 5.09917638 | 0.47929085 | 3463.08966 | 559.287931 |
| GAK9 | 128 | 79.5313558 | 92.2176175 | 85.8744867 | 8.9705417 | 24.6461394 | 17.1731739 | 12.2426331 | 2.09260336 | -25.452 | 0.24890159 | 5.0965 | 1.46724657 | 7.01438046 | 0.20032061 | 2043.97116 | 155.585334 |
| GAK9 | 103 | 90.7505832 | 105.302244 | 98.0264137 | 10.2895781 | 5.88837187 | 8.28478311 | 13.79215 | 0.02288467 | -25.517 | 0.22485996 | 7.134 | 0.34648232 | 7.10740629 | 0.10498052 | 2055.55173 | 171.962737 |
| GAK9 | 78 | 106.781303 | 122.111871 | 114.446587 | 10.8403488 | 36.4651237 | -8.11258402 | 16.3767931 | 2.45714824 | -25.735 | 0.0212132 | 6.9725 | 0.07848885 | 6.98833929 | 0.17743549 | 1852.8917 | 409.435089 |
| GAK9 | 53 | 119.095273 | 168.667022 | 143.881148 | 35.0525195 | 2.33906648 | -33.0679009 | 22.0583155 | 6.01487688 | -25.722 | 0.22627417 | 7.0375 | 0.06434672 | 6.52276225 | 0.36565848 | 1742.87625 | 155.585334 |

| | | | | | | | | | | | | | | | | | |
|---|---|---|---|---|---|---|---|---|---|---|---|---|---|---|---|---|---|
| GAK9 | 27 | 233.729753 | 406.737059 | 320.233406 | 122.334639 | 83.8880951 | -9.09045155 | 75.0201845 | 38.4287259 | -25.8755 | 0.1039447 | 7.547 | 0.15697771 | 4.26862995 | 0.63900874 | 3323.62448 | 720.605757 |
| GAK5 | 103 | 177.068312 | 235.802552 | 206.435432 | 41.5313796 | -57.1092458 | 13.5473529 | 26.5453709 | 0.66102355 | -26.476 | 0.58689863 | 7.524 | 0.04384062 | 7.77670173 | 0.20271864 | 6863.58053 | 4290.46649 |
| GAK5 | 78 | 139.427004 | 232.319631 | 185.873317 | 65.685006 | 54.3422085 | 126.641317 | 30.3942415 | 16.8070252 | -25.7475 | 0.16334167 | 10.938 | 2.81428499 | 6.11541228 | 0.65624275 | 5421.65184 | 326.271216 |
| GAK5 | 53 | 148.019744 | 172.61403 | 160.316887 | 17.3907863 | 19.9196281 | 35.877873 | 27.0795028 | 3.49568638 | -25.8585 | 0.15202796 | 5.671 | 0.89943983 | 5.9202301 | 0.16861657 | 3945.11687 | 1076.69501 |
| GAK5 | 27 | 333.54806 | 385.592982 | 359.570521 | 36.8013172 | 128.660529 | 140.955857 | 80.1854566 | 6.1803375 | -25.219 | 0.30547013 | 8.12 | 0.41295036 | 4.48423612 | 0.12812396 | 7071.21826 | 962.500086 |
| GAK1 | 128 | 420.780138 | 509.823713 | 465.301925 | 62.9633155 | -172.371592 | -46.5103511 | 41.9196491 | 3.26700873 | -25.5775 | 0.62861793 | -6.4005 | 4.00010306 | 11.0998526 | 0.15615566 | 11714.503 | 1277.09876 |
| GAK1 | 103 | 204.324004 | 215.165106 | 209.744555 | 7.66581686 | 32.1764672 | -6.13545288 | 37.7091452 | 3.72263822 | -24.192 | 0.17253405 | 6.0775 | 2.93661446 | 5.5621668 | 0.10526811 | 9607.39751 | 1241.6238 |
| GAK1 | 78 | 280.251202 | 331.376429 | 305.813816 | 36.1509952 | 13.2956731 | -24.274259 | 52.3910747 | 1.22403781 | -24.354 | 0.50911688 | 2.1425 | 1.53230039 | 5.83713576 | 0.12049909 | 10209.4276 | 957.824074 |
| GAK1 | 53 | 300.469359 | 416.069757 | 358.269558 | 81.7418255 | 116.832782 | 6.9802939 | 58.7192958 | 14.4653551 | -23.653 | 0.08909545 | 4.962 | 1.12995664 | 6.10139399 | 0.33577207 | 9645.02439 | 2997.6346 |
| GAK1 | 27 | 330.432128 | 465.869331 | 398.150729 | 95.7685648 | 341.7592 | 519.382827 | 85.3605469 | 22.7929913 | -23.982 | 0.07071068 | 6.8865 | 1.59028315 | 4.66434136 | 0.35938298 | 10610.7811 | 1986.59808 |
| KOD5 | 53 | 616.771642 | 650.017259 | 633.394451 | 23.5082017 | 257.542478 | -13.6566844 | 110.387519 | 2.88401831 | -22.0345 | 0.04879037 | 7.0055 | 0.95105862 | 5.73791724 | 0.0453881 | 23891.2799 | 459.841098 |
| KOD5 | 27 | 979.97741 | 1097.10914 | 1038.54327 | 82.8246387 | -503.286745 | -60.2499186 | 268.115741 | 23.5213043 | -20.8035 | 0.08131728 | 6.9815 | 0.42072853 | 3.87348863 | 0.11855977 | 16087.5177 | 810.19622 |
| KOD10 | 128 | 77.1346214 | 81.5740476 | 79.3543345 | 3.13914843 | 6.25500155 | 14.0825056 | 11.7435612 | 0.59077013 | -25.716 | 0.11737973 | 19.2305 | 15.2869415 | 6.75726324 | 0.06399661 | 1566.84633 | 250.177215 |
| KOD10 | 103 | 83.701317 | 121.012642 | 102.356979 | 26.3830908 | -0.74309587 | -16.9137419 | 16.7283715 | 6.37254116 | -25.8275 | 0.17324116 | 8.543 | 1.62351717 | 6.11876533 | 0.45995095 | 2343.95157 | 205.502712 |
| KOD10 | 78 | 131.53207 | 139.13821 | 135.33514 | 5.37835351 | 65.3146769 | 37.2921704 | 25.7993575 | 0.35101921 | -25.7315 | 0.25950819 | 5.023 | 1.50472323 | 5.24567871 | 0.04200551 | 1339.40089 | 411.005424 |
| KOD10 | 53 | 384.027526 | 579.892342 | 481.959934 | 138.49734 | 124.17041 | -62.5027174 | 122.993358 | 39.6464108 | -25.7235 | 0.03606245 | 8.0485 | 0.60316208 | 3.91858506 | 0.43183825 | 4864.80514 | 1483.19349 |
| KOD10 | 27 | 1971.48593 | 2289.37332 | 2130.42962 | 224.780331 | 1643.53337 | 625.657804 | 1240.58179 | 176.22146 | -29.056 | 0.83580022 | 9.5865 | 1.28764145 | 1.71728268 | 0.17694549 | 44509.808 | 8389.87159 |

Supporting Information Table 4.1. Empirical Carbon to Volume values A and B for different particle types Mini Literature Review

| A Low | A Low Uncertainty | A High | A High Uncertainty | A Units | B | B Uncertainty | Particle Type | Other Information | Source |
|---|---|---|---|---|---|---|---|---|---|
| 1.00E-10 | 4.00E-11 | | | mg C μm−3 | 1 | | Euphausiid FP | Minimization (gel to trap) | (Durkin et al. 2021) |
| 1.00E-10 | 4.00E-11 | | | mg C μm−3 | 1 | | Copepod FP | Minimization (gel to trap) | (Durkin et al. 2021) |

| | | | | | | | | | |
|---|---|---|---|---|---|---|---|---|---|
| 1.00E-10 | 4.00E-11 | | | mg C µm−3 | 1 | | Round mini FP | Minimization (gel to trap) | (Durkin et al. 2021) |
| 1.00E-10 | 4.00E-11 | | | mg C µm−3 | 0.8 | | Aggregate | Minimization (gel to trap) | (Durkin et al. 2021) |
| 1.00E-10 | 4.00E-11 | | | mg C µm−3 | 0.83 | | Dense detritus | Minimization (gel to trap) | (Durkin et al. 2021) |
| 1.00E-10 | 4.00E-11 | | | mg C µm−3 | 0.83 | | Large loose FP | Minimization (gel to trap) | (Durkin et al. 2021) |
| 4.20E-11 | | | | mg C µm−3 | 1 | | Larvacean FP | Empirical from trap, nets, and incubation | (Gonzalez et al. 1994) |
| 5.70E-11 | | | | mg C µm−3 | 1 | | Copepod FP | Empirical from trap, nets, and incubation | (Gonzalez et al. 1994) |
| 1.60E-11 | | | | mg C µm−3 | 1 | | Euphausiid FP | Empirical from trap, nets, and incubation | (Gonzalez et al. 1994) |
| 1.80E-10 | 7.00E-11 | | | mg C µm−3 | 1 | | Copepod FP | Empirical from trap (fixed) | (Lundsgaard and Olesen 1997) |
| 1.20E-10 | 6.00E-11 | | | mg C µm−3 | 1 | | Copepod FP | Empirical from trap (not fixed) | (Lundsgaard and Olesen 1997) |
| 1.80E-10 | 7.00E-11 | | | mg C µm−3 | 1 | | Copepod FP | Empirical from incubation (not fixed) | (Lundsgaard and Olesen 1997) |
| 3.00E-10 | 7.00E-11 | | | mg C µm−3 | 1 | | Copepod FP | Empirical from incubation (fixed) | (Lundsgaard and Olesen 1997) |
| 7.80E-11 | 3.20E-11 | | | mg C µm−3 | 1 | | Copepod FP | Empirical from incubation (not fixed) | (Lundsgaard and Olesen 1997) |
| 6.10E-11 | 1.00E-11 | | | mg C µm−3 | 1 | | Copepod FP | Empirical from incubation (fixed) | (Lundsgaard and Olesen 1997) |
| 3.80E-10 | | 7.50E-10 | | mg C µm−3 | 1 | | Copepod FP | Empirical natural seston | (Honjo and Roman 1978) |
| 2.00E-12 | | 1.60E-11 | | mg C µm−3 | 1 | | Salp FP | Empirical natural seston | (Madin 1982) |

| | | | | | | | | | |
|---|---|---|---|---|---|---|---|---|---|
| 5.00E-12 | | 6.00E-12 | | mg C µm−3 | 1 | | Salp FP | Empirical natural seston | (Madin 1982) |
| 2.00E-12 | | 7.00E-12 | | mg C µm−3 | 1 | | Salp FP | Empirical natural seston | (Madin 1982) |
| 2.10E-10 | | | | mg C µm−3 | 1 | | Copepod FP | Empirical from incubation (fed) | (Abou Debs 1984) |
| 1.00E-11 | | | | mg C µm−3 | 1 | | Salp FP | Empirical natural seston | (Caron et al. 1989) |
| 1.94E-09 | | | | mg C µm−3 | 1 | | Salp FP | Empirical natural seston | (Caron et al. 1989) |
| 1.00E-11 | | 2.00E-11 | | mg C µm−3 | 1 | | Euphausiid FP | Empirical natural seston | (Youngbluth et al. 1989) |
| 6.86E-09 | | 5.07E-07 | | mg C µm−3 | 1 | | Copepod FP | Empirical natural seston | (Lane et al. 1994) |
| 1.70E-10 | | 2.50E-09 | | mg C µm−3 | 1 | | Copepod FP | Empirical from incubation (fed) | (Butler and Dam 1994) |
| 2.80E-10 | | | | mg C µm−3 | 1 | | Copepod FP | Empirical from incubation (fed) | (Hansen et al. 1996) |
| 3.90E-10 | | | | mg C µm−3 | 1 | | Copepod FP | Empirical from incubation (fed) | (Hansen et al. 1996) |
| 1.00E-11 | | 6.00E-11 | | mg C µm−3 | 1 | | Copepod FP | Empirical natural seston | (Urban-Rich 1997) |
| 1.00E-11 | | 2.50E-10 | | mg C µm−3 | 1 | | Copepod FP | Empirical natural seston | (Urban-Rich et al. 1998) |
| 3.50E-11 | | | | mg C µm−3 | 1 | | Salp FP | Empirical from incubation | (Silver and Bruland 1981) |
| 2.01E-11 | | | | mg C µm−3 | 1 | | Pteropod FP | Empirical from incubation | (Silver and Bruland 1981) |
| 1.10E-12 | | | | mg C µm−3 | 1 | | Rhizaria | Volume calculated, carbon from Menden-Deuer and Lessard, 2000 | (Stukel et al. 2018) |

| | | | | | | | | | |
|---|---|---|---|---|---|---|---|---|---|
| 1.09E-12 | 2.90E-13 | | | mg C µm−3 | 0.39 | 0.08 | Larvacean House | Empirical natural seston | (Alldredge 1998) |
| 9.70E-13 | 2.40E-13 | | | mg C µm−3 | 0.5 | 0.06 | Diatom | Empirical natural seston | (Alldredge 1998) |
| 1.05E-12 | 1.60E-13 | | | mg C µm−3 | 0.51 | 0.05 | Fecal Pellet | Empirical natural seston | (Alldredge 1998) |
| 9.90E-13 | 1.20E-13 | | | mg C µm−3 | 0.52 | 0.03 | All Types | Empirical natural seston | (Alldredge 1998) |
| 2.88E-10 | | | | mg C µm−3 | 0.88 | 0.028 | Diatom | Empirical grown in culture | (Menden-Deuer and Lessard 2000) |
| 2.15E-10 | | | | mg C µm−3 | 0.94 | 0.041 | Rhizaria | Empirical grown in culture | (Menden-Deuer and Lessard 2000) |
| 2.00E-11 | | 3.90E-10 | | mg C µm−3 | 1 | | Copepod FP | Empirical from incubation (fed) | (Olesen et al. 2005; Turner 2015) |
| 3.50E-11 | | | | mg C µm−3 | 1 | | Round mini FP | Empirical from trap | (Manno et al. 2015; Belcher et al. 2016) |
| 5.20E-11 | | | | mg C µm−3 | 1 | | Copepod FP | Empirical from trap | (Manno et al. 2015; Belcher et al. 2016) |
| 3.00E-11 | | | | mg C µm−3 | 1 | | Euphausiid FP | Empirical from trap | (Manno et al. 2015; Belcher et al. 2016) |
| 2.77E-11 | 1.08E-11 | | | mg C µm−3 | 1 | | Salp FP | Empirical from incubation | (Iversen et al. 2017) |
| 2.00E-11 | | | | mg C µm−3 | 1 | | All Types | Empirical from trap | (Gleiber et al. 2012) |
| 1.60E-11 | 2.00E-12 | | | mg C µm−3 | 1 | | Euphausiid FP | Empirical from trap | (Gleiber et al. 2012) |
| 4.10E-11 | 1.90E-11 | | | mg C µm−3 | 1 | | Copepod FP | Empirical from trap | (Gleiber et al. 2012) |
| 2.10E-11 | 2.10E-11 | | | mg C µm−3 | 1 | | Salp FP | Empirical from trap | (Gleiber et al. 2012) |
| 1.00E-11 | | 1.50E-10 | | mg C µm−3 | 1 | | Fecal Pellet | Review Range | (Urrère and Knauer 1981; Gonzalez 1992; Wassmann et al. 2000; Wilson et al. 2008; Wexels Riser et al. 2008; |

| | | | | | | | | | |
|---|---|---|---|---|---|---|---|---|---|
| | | | | | | | | | Smith et al. 2011; Gleiber et al. 2012) |
| 1.30E-10 | 4.00E-11 | | | mg C µm−3 | 1 | | Copepod FP | Empirical from incubation | (Wilson et al. 2008) |
| 1.50E-10 | 3.00E-11 | | | mg C µm−3 | 1 | | Copepod FP | Empirical from incubation | (Wilson et al. 2008) |
| 8.00E-11 | 1.00E-11 | | | mg C µm−3 | 1 | | Euphausiid FP | Empirical from incubation | (Wilson et al. 2008) |
| 3.00E-11 | 1.00E-11 | | | mg C µm−3 | 1 | | Chaetognath FP | Empirical from incubation | (Wilson et al. 2008) |
| 1.00E-11 | | 1.50E-10 | | mg C µm−3 | 1 | | Fecal Pellet | Review Range | (Silver and Gowing 1991; Lundsgaard and Olesen 1997; Carroll et al. 1998; Taguchi and Saino 1998; Urban-Rich et al. 1998; Roy et al. 2000; Wassmann et al. 2000; Gowing et al. 2001; Wexels Riser et al. 2001; Suzuki et al. 2003; Huskin et al. 2004; Reigstad et al. 2005; Olesen et al. 2005; Wilson et al. 2008) |
| 8.00E-11 | 4.40E-11 | | | mg C µm−3 | 1 | | Euphausiid FP | Empirical from incubation | (Pauli et al. 2021) |
| 2.60E-11 | 1.20E-11 | | | mg C µm−3 | 1 | | Salp FP | Empirical from incubation | (Pauli et al. 2021) |
| 1.70E-11 | 8.00E-12 | | | mg C µm−3 | 1 | | Salp FP | Empirical from incubation | (Pauli et al. 2021) |

| | | | | | | | | | |
|---|---|---|---|---|---|---|---|---|---|
| 2.00E-11 | | 6.00E-11 | | mg C µm−3 | 1 | | Euphausiid FP | Review Range | (Gleiber et al. 2012; Belcher et al. 2016; Pauli et al. 2021) |

Supporting Information Table 4.2. Empirical Carbon to Volume values A and B for different particle types Mini Literature Review Part 2

| **Original A Low** | **Original A Low Uncertainty** | **Original A High** | **Original A High Uncertainty** | **Original A Units** | **A low in mg C µm−3** | **A high in mg C µm−3** | **Original B** | **Original B Uncertainty** | **Notes** |
|---|---|---|---|---|---|---|---|---|---|
| 1.00E-10 | | | | mg C µm−3 | 1.00E-10 | | 1 | | |
| 1.00E-10 | | | | mg C µm−3 | 1.00E-10 | | 1 | | |
| 1.00E-10 | | | | mg C µm−3 | 1.00E-10 | | 1 | | |
| 1.00E-10 | | | | mg C µm−3 | 1.00E-10 | | 0.8 | | |
| 1.00E-10 | | | | mg C µm−3 | 1.00E-10 | | 0.83 | | |
| 1.00E-10 | | | | mg C µm−3 | 1.00E-10 | | 0.83 | | |
| 4.20E-02 | | | | mg C mm−3 | 4.20E-11 | | | | |
| 5.70E-02 | | | | mg C mm−3 | 5.70E-11 | | | | |
| 1.60E-02 | | | | mg C mm−3 | 1.60E-11 | | | | |

| | | | | | | | | | |
|---|---|---|---|---|---|---|---|---|---|
| 1.80E-01 | 7.00E-02 | | | pg C μm−3 | 1.80E-10 | | | | |
| 1.20E-01 | 6.00E-02 | | | pg C μm−3 | 1.20E-10 | | | | |
| 1.80E-01 | 7.00E-02 | | | pg C μm−3 | 1.80E-10 | | | | |
| 3.00E-01 | 7.00E-02 | | | pg C μm−3 | 3.00E-10 | | | | |
| 7.80E-02 | 3.20E-02 | | | pg C μm−3 | 7.80E-11 | | | | |
| 6.10E-02 | 1.00E-02 | | | pg C μm−3 | 6.10E-11 | | | | |
| 3.80E-01 | | 7.50E-01 | | pg C μm−3 | 3.80E-10 | 7.50E-10 | | | * |
| 2.00E-03 | | 1.60E-02 | | pg C μm−3 | 2.00E-12 | 1.60E-11 | | | * |
| 5.00E-03 | | 6.00E-03 | | pg C μm−3 | 5.00E-12 | 6.00E-12 | | | * |
| 2.00E-03 | | 7.00E-03 | | pg C μm−3 | 2.00E-12 | 7.00E-12 | | | * |
| 2.10E-01 | | | | pg C μm−3 | 2.10E-10 | | | | * |
| 1.00E-02 | | | | pg C μm−3 | 1.00E-11 | | | | * |
| 1.94E+00 | | | | pg C μm−3 | 1.94E-09 | | | | * |
| 1.00E-02 | | 2.00E-02 | | pg C μm−3 | 1.00E-11 | 2.00E-11 | | | * |
| 6.86E+00 | | ####### | | pg C μm−3 | 6.86E-09 | 5.07E-07 | | | * |
| 1.70E-01 | | ####### | | pg C μm−3 | 1.70E-10 | 2.50E-09 | | | * |
| 2.80E-01 | | | | pg C μm−3 | 2.80E-10 | | | | * |
| 3.90E-01 | | | | pg C μm−3 | 3.90E-10 | | | | * |
| 1.00E-02 | | 6.00E-02 | | pg C μm−3 | 1.00E-11 | 6.00E-11 | | | * |

| | | | | | | | | | |
|---|---|---|---|---|---|---|---|---|---|
| 1.00E-02 | | 2.50E-01 | | pg C μm−3 | 1.00E-11 | 2.50E-10 | | | * |
| 3.50E-08 | | | | μg C μm−3 | 3.50E-11 | | | | |
| 2.01E-08 | | | | μg C μm−3 | 2.01E-11 | | | | |
| 1.10E+00 | | | | μg C mm−3 | 1.10E-12 | | | | |
| 1.09E+00 | 2.90E-01 | | | μg C mm−3 | 1.09E-12 | | 0.39 | 0.08 | |
| 9.70E-01 | 2.40E-01 | | | μg C mm−3 | 9.70E-13 | | 0.56 | 0.06 | |
| 1.05E+00 | 1.60E-01 | | | μg C mm−3 | 1.05E-12 | | 0.51 | 0.05 | |
| 9.90E-01 | 1.20E-01 | | | μg C mm−3 | 9.90E-13 | | 0.52 | 0.03 | |
| -5.41E-01 | 9.90E-02 | | | log(pg C μm−3) | 2.88E-10 | | | | |
| -6.65E-01 | 1.32E-01 | | | log(pg C μm−3) | 2.16E-10 | | | | |
| 2.00E-02 | | 3.90E-01 | | pg C μm−3 | 2.00E-11 | 3.90E-10 | | | |
| 3.50E-02 | | | | mg C mm-3 | 3.50E-11 | | | | |
| 5.20E-02 | | | | mg C mm-3 | 5.20E-11 | | | | |
| 3.00E-02 | | | | mg C mm-3 | 3.00E-11 | | | | |
| 2.77E+01 | 1.08E+01 | | | μg C mm−3 | 2.77E-11 | | | | |
| 2.00E-02 | | | | mg C mm-3 | 2.00E-11 | | | | |
| 1.60E-02 | 2.00E-03 | | | mg C mm-3 | 1.60E-11 | | | | |

| | | | | | | |
|---|---|---|---|---|---|---|
| 4.10E-02 | 1.90E-02 | | | mg C mm-3 | 4.10E-11 | |
| 2.10E-02 | 2.10E-02 | | | mg C mm-3 | 2.10E-11 | |
| 1.00E-02 | | 1.50E-01 | | mg C mm-3 | 1.00E-11 | 1.50E-10 |
| 1.30E-01 | 4.00E-02 | | | mg C mm-3 | 1.30E-10 | |
| 1.50E-01 | 3.00E-02 | | | mg C mm-3 | 1.50E-10 | |
| 8.00E-02 | 1.00E-02 | | | mg C mm-3 | 8.00E-11 | |
| 3.00E-02 | 1.00E-02 | | | mg C mm-3 | 3.00E-11 | |
| 1.00E-02 | | 1.50E-01 | | mg C mm-3 | 1.00E-11 | 1.50E-10 |
| 8.00E-02 | 4.40E-02 | | | mg C mm-3 | 8.00E-11 | |
| 2.60E-02 | 1.20E-02 | | | mg C mm-3 | 2.60E-11 | |
| 1.70E-02 | 8.00E-03 | | | mg C mm-3 | 1.70E-11 | |
| 2.00E-02 | | 6.00E-02 | | mg C mm-3 | 2.00E-11 | 6.00E-11 |

* Zooplankton fecal pellet carbon concentrations as reported in the literature and scaled by an average volume

Supporting Information Table 5. Gel-derived carbon flux total (est_c_flux in mg C $m^{-2}$ $d^{-1}$) and volume flux (est_v_flux in $\mu m^{-3}$) by type (C_Flux in mg C $m^{-2}$ $d^{-1}$ and Percent_C_Flux) for each tube along with chemically derived carbon flux (trap_c_flux in mg C $m^{-2}$ $d^{-1}$) and uncertainty.

| station | depth | tube | trap_c_flux | trap_c_flux uncertainty | est_c_flux | est_v_flux | Type | Percent_C_Flux | C_Flux |
|---|---|---|---|---|---|---|---|---|---|
| GAK1 | 103 | B1 | 209.74455527955 | 7.66581685911836 | 698.8356585 | 165504855545404.34 | aggregate | 0.674754311 | 471.5423735 |

| | | | | | | | | | |
|---|---|---|---|---|---|---|---|---|---|
| GAK1 | 103 | B1 | 209.74455527955 | 7.66581685911836 | 698.8356585 | 165504855545404.34 | dense_detritus | 0.003447909 | 2.409521779 |
| GAK1 | 103 | B1 | 209.74455527955 | 7.66581685911836 | 698.8356585 | 165504855545404.34 | unidentified | 0.037326523 | 26.08510521 |
| GAK1 | 103 | B1 | 209.74455527955 | 7.66581685911836 | 698.8356585 | 165504855545404.34 | large_loose_pellet | 0.014919125 | 10.42601661 |
| GAK1 | 103 | B1 | 209.74455527955 | 7.66581685911836 | 698.8356585 | 165504855545404.34 | long_fecal_pellet | 0.237761354 | 166.1561122 |
| GAK1 | 103 | B1 | 209.74455527955 | 7.66581685911836 | 698.8356585 | 165504855545404.34 | mini_pellet | 0.021296237 | 14.88256971 |
| GAK1 | 103 | B1 | 209.74455527955 | 7.66581685911836 | 698.8356585 | 165504855545404.34 | salp_pellet | 0 | 0 |
| GAK1 | 103 | B1 | 209.74455527955 | 7.66581685911836 | 698.8356585 | 165504855545404.34 | short_pellet | 0.007839947 | 5.47883444 |
| GAK1 | 103 | B1 | 209.74455527955 | 7.66581685911836 | 698.8356585 | 165504855545404.34 | phytoplankton | 0.000781014 | 0.54580025 |
| GAK1 | 103 | B1 | 209.74455527955 | 7.66581685911836 | 698.8356585 | 165504855545404.34 | rhizaria | 0.00187358 | 1.30932484 |
| GAK1 | 128 | A1 | 465.301925379784 | 62.9633155279178 | 398.4386928 | 73616185419736.6 | aggregate | 0.65222518 | 259.8717481 |
| GAK1 | 128 | A1 | 465.301925379784 | 62.9633155279178 | 398.4386928 | 73616185419736.6 | dense_detritus | 0.005859691 | 2.334727682 |
| GAK1 | 128 | A1 | 465.301925379784 | 62.9633155279178 | 398.4386928 | 73616185419736.6 | unidentified | 0.034359637 | 13.69020879 |
| GAK1 | 128 | A1 | 465.301925379784 | 62.9633155279178 | 398.4386928 | 73616185419736.6 | large_loose_pellet | 0.046047451 | 18.34708601 |
| GAK1 | 128 | A1 | 465.301925379784 | 62.9633155279178 | 398.4386928 | 73616185419736.6 | long_fecal_pellet | 0.253263098 | 100.9098177 |
| GAK1 | 128 | A1 | 465.301925379784 | 62.9633155279178 | 398.4386928 | 73616185419736.6 | mini_pellet | 0.00146583 | 0.584043357 |
| GAK1 | 128 | A1 | 465.301925379784 | 62.9633155279178 | 398.4386928 | 73616185419736.6 | salp_pellet | 0.000598395 | 0.238423682 |
| GAK1 | 128 | A1 | 465.301925379784 | 62.9633155279178 | 398.4386928 | 73616185419736.6 | short_pellet | 0.004724377 | 1.882374716 |
| GAK1 | 128 | A1 | 465.301925379784 | 62.9633155279178 | 398.4386928 | 73616185419736.6 | phytoplankton | 0.000308202 | 0.12279977 |
| GAK1 | 128 | A1 | 465.301925379784 | 62.9633155279178 | 398.4386928 | 73616185419736.6 | rhizaria | 0.001148139 | 0.457462911 |
| GAK1 | 27 | E1 | 398.150729227848 | 95.7685648040826 | 369.6176719 | 86202456022977.02 | aggregate | 0.84124126 | 310.9376362 |
| GAK1 | 27 | E1 | 398.150729227848 | 95.7685648040826 | 369.6176719 | 86202456022977.02 | dense_detritus | 0.008226947 | 3.040825008 |
| GAK1 | 27 | E1 | 398.150729227848 | 95.7685648040826 | 369.6176719 | 86202456022977.02 | unidentified | 0.04112505 | 15.20054519 |
| GAK1 | 27 | E1 | 398.150729227848 | 95.7685648040826 | 369.6176719 | 86202456022977.02 | large_loose_pellet | 0.013529816 | 5.000859232 |
| GAK1 | 27 | E1 | 398.150729227848 | 95.7685648040826 | 369.6176719 | 86202456022977.02 | long_fecal_pellet | 0.083038329 | 30.69243402 |
| GAK1 | 27 | E1 | 398.150729227848 | 95.7685648040826 | 369.6176719 | 86202456022977.02 | mini_pellet | 0.001351784 | 0.499643158 |
| GAK1 | 27 | E1 | 398.150729227848 | 95.7685648040826 | 369.6176719 | 86202456022977.02 | salp_pellet | 0 | 0 |
| GAK1 | 27 | E1 | 398.150729227848 | 95.7685648040826 | 369.6176719 | 86202456022977.02 | short_pellet | 0.006954502 | 2.570506791 |
| GAK1 | 27 | E1 | 398.150729227848 | 95.7685648040826 | 369.6176719 | 86202456022977.02 | phytoplankton | 0.002336854 | 0.863742698 |
| GAK1 | 27 | E1 | 398.150729227848 | 95.7685648040826 | 369.6176719 | 86202456022977.02 | rhizaria | 0.002195457 | 0.81147961 |
| GAK1 | 53 | D1 | 358.269558207252 | 81.7418255148796 | 618.2689654 | 151269580189269.12 | aggregate | 0.669937113 | 414.201326 |
| GAK1 | 53 | D1 | 358.269558207252 | 81.7418255148796 | 618.2689654 | 151269580189269.12 | dense_detritus | 0.010159969 | 6.281593447 |

| | | | | | | | | | |
|---|---|---|---|---|---|---|---|---|---|
| GAK1 | 53 | D1 | 358.269558207252 | 81.7418255148796 | 618.2689654 | 151269580189269.12 | unidentified | 0.026949981 | 16.66233714 |
| GAK1 | 53 | D1 | 358.269558207252 | 81.7418255148796 | 618.2689654 | 151269580189269.12 | large_loose_pellet | 0.023699233 | 14.65250002 |
| GAK1 | 53 | D1 | 358.269558207252 | 81.7418255148796 | 618.2689654 | 151269580189269.12 | long_fecal_pellet | 0.255505 | 157.9708122 |
| GAK1 | 53 | D1 | 358.269558207252 | 81.7418255148796 | 618.2689654 | 151269580189269.12 | mini_pellet | 0.000762929 | 0.471695091 |
| GAK1 | 53 | D1 | 358.269558207252 | 81.7418255148796 | 618.2689654 | 151269580189269.12 | salp_pellet | 0.002657825 | 1.643250803 |
| GAK1 | 53 | D1 | 358.269558207252 | 81.7418255148796 | 618.2689654 | 151269580189269.12 | short_pellet | 0.009460093 | 5.848882093 |
| GAK1 | 53 | D1 | 358.269558207252 | 81.7418255148796 | 618.2689654 | 151269580189269.12 | phytoplankton | 0.000625719 | 0.386862522 |
| GAK1 | 53 | D1 | 358.269558207252 | 81.7418255148796 | 618.2689654 | 151269580189269.12 | rhizaria | 0.000242138 | 0.149706122 |
| GAK1 | 78 | C3 | 305.813815570576 | 36.1509952351086 | 467.9272387 | 89258383880336.69 | aggregate | 0.680798857 | 318.5643293 |
| GAK1 | 78 | C3 | 305.813815570576 | 36.1509952351086 | 467.9272387 | 89258383880336.69 | dense_detritus | 0.012338421 | 5.773483158 |
| GAK1 | 78 | C3 | 305.813815570576 | 36.1509952351086 | 467.9272387 | 89258383880336.69 | unidentified | 0.04379277 | 20.49182987 |
| GAK1 | 78 | C3 | 305.813815570576 | 36.1509952351086 | 467.9272387 | 89258383880336.69 | large_loose_pellet | 0.011932996 | 5.583773773 |
| GAK1 | 78 | C3 | 305.813815570576 | 36.1509952351086 | 467.9272387 | 89258383880336.69 | long_fecal_pellet | 0.241497366 | 113.0031955 |
| GAK1 | 78 | C3 | 305.813815570576 | 36.1509952351086 | 467.9272387 | 89258383880336.69 | mini_pellet | 0.000209743 | 0.098144663 |
| GAK1 | 78 | C3 | 305.813815570576 | 36.1509952351086 | 467.9272387 | 89258383880336.69 | salp_pellet | 0 | 0 |
| GAK1 | 78 | C3 | 305.813815570576 | 36.1509952351086 | 467.9272387 | 89258383880336.69 | short_pellet | 0.007816435 | 3.657523066 |
| GAK1 | 78 | C3 | 305.813815570576 | 36.1509952351086 | 467.9272387 | 89258383880336.69 | phytoplankton | 0.000839359 | 0.392759027 |
| GAK1 | 78 | C3 | 305.813815570576 | 36.1509952351086 | 467.9272387 | 89258383880336.69 | rhizaria | 0.000774053 | 0.362200372 |
| GAK15 | 103 | B1 | 79.6793405026835 | 7.69514011163473 | 159.8535201 | 32439502565871.785 | aggregate | 0.439831343 | 70.30858837 |
| GAK15 | 103 | B1 | 79.6793405026835 | 7.69514011163473 | 159.8535201 | 32439502565871.785 | dense_detritus | 0 | 0 |
| GAK15 | 103 | B1 | 79.6793405026835 | 7.69514011163473 | 159.8535201 | 32439502565871.785 | unidentified | 0.282501169 | 45.15880634 |
| GAK15 | 103 | B1 | 79.6793405026835 | 7.69514011163473 | 159.8535201 | 32439502565871.785 | large_loose_pellet | 0.001382781 | 0.221042383 |
| GAK15 | 103 | B1 | 79.6793405026835 | 7.69514011163473 | 159.8535201 | 32439502565871.785 | long_fecal_pellet | 0.155971753 | 24.93263371 |
| GAK15 | 103 | B1 | 79.6793405026835 | 7.69514011163473 | 159.8535201 | 32439502565871.785 | mini_pellet | 0.078839806 | 12.60282053 |
| GAK15 | 103 | B1 | 79.6793405026835 | 7.69514011163473 | 159.8535201 | 32439502565871.785 | salp_pellet | 0 | 0 |
| GAK15 | 103 | B1 | 79.6793405026835 | 7.69514011163473 | 159.8535201 | 32439502565871.785 | short_pellet | 0.040393611 | 6.457060931 |
| GAK15 | 103 | B1 | 79.6793405026835 | 7.69514011163473 | 159.8535201 | 32439502565871.785 | phytoplankton | 0 | 0 |
| GAK15 | 103 | B1 | 79.6793405026835 | 7.69514011163473 | 159.8535201 | 32439502565871.785 | rhizaria | 0.001079537 | 0.172567844 |
| GAK15 | 103 | B3 | 79.6793405026835 | 7.69514011163473 | 144.6205087 | 34759477369743.504 | aggregate | 0.512315579 | 74.09133956 |
| GAK15 | 103 | B3 | 79.6793405026835 | 7.69514011163473 | 144.6205087 | 34759477369743.504 | dense_detritus | 0.011956963 | 1.72922206 |
| GAK15 | 103 | B3 | 79.6793405026835 | 7.69514011163473 | 144.6205087 | 34759477369743.504 | unidentified | 0.189865263 | 27.45841088 |

| | | | | | | | | | |
|---|---|---|---|---|---|---|---|---|---|
| GAK15 | 103 | B3 | 79.6793405026835 | 7.69514011163473 | 144.6205087 | 34759477369743.504 | large_loose_pellet | 0 | 0 |
| GAK15 | 103 | B3 | 79.6793405026835 | 7.69514011163473 | 144.6205087 | 34759477369743.504 | long_fecal_pellet | 0.138348449 | 20.008023 |
| GAK15 | 103 | B3 | 79.6793405026835 | 7.69514011163473 | 144.6205087 | 34759477369743.504 | mini_pellet | 0.061421849 | 8.882859067 |
| GAK15 | 103 | B3 | 79.6793405026835 | 7.69514011163473 | 144.6205087 | 34759477369743.504 | salp_pellet | 0 | 0 |
| GAK15 | 103 | B3 | 79.6793405026835 | 7.69514011163473 | 144.6205087 | 34759477369743.504 | short_pellet | 0.071787264 | 10.38191062 |
| GAK15 | 103 | B3 | 79.6793405026835 | 7.69514011163473 | 144.6205087 | 34759477369743.504 | phytoplankton | 0.002314389 | 0.334708168 |
| GAK15 | 103 | B3 | 79.6793405026835 | 7.69514011163473 | 144.6205087 | 34759477369743.504 | rhizaria | 0.011990245 | 1.734035307 |
| GAK15 | 128 | A1 | 72.3390400151472 | 5.28551875731876 | 49.26522008 | 13827113979658.605 | aggregate | 0.411164833 | 20.25612599 |
| GAK15 | 128 | A1 | 72.3390400151472 | 5.28551875731876 | 49.26522008 | 13827113979658.605 | dense_detritus | 0 | 0 |
| GAK15 | 128 | A1 | 72.3390400151472 | 5.28551875731876 | 49.26522008 | 13827113979658.605 | unidentified | 0.496144178 | 24.44265211 |
| GAK15 | 128 | A1 | 72.3390400151472 | 5.28551875731876 | 49.26522008 | 13827113979658.605 | large_loose_pellet | 0 | 0 |
| GAK15 | 128 | A1 | 72.3390400151472 | 5.28551875731876 | 49.26522008 | 13827113979658.605 | long_fecal_pellet | 0.009216165 | 0.454036374 |
| GAK15 | 128 | A1 | 72.3390400151472 | 5.28551875731876 | 49.26522008 | 13827113979658.605 | mini_pellet | 0.036332864 | 1.789946546 |
| GAK15 | 128 | A1 | 72.3390400151472 | 5.28551875731876 | 49.26522008 | 13827113979658.605 | salp_pellet | 0 | 0 |
| GAK15 | 128 | A1 | 72.3390400151472 | 5.28551875731876 | 49.26522008 | 13827113979658.605 | short_pellet | 0.045832583 | 2.257952283 |
| GAK15 | 128 | A1 | 72.3390400151472 | 5.28551875731876 | 49.26522008 | 13827113979658.605 | phytoplankton | 0 | 0 |
| GAK15 | 128 | A1 | 72.3390400151472 | 5.28551875731876 | 49.26522008 | 13827113979658.605 | rhizaria | 0.001309378 | 0.064506781 |
| GAK15 | 128 | A3 | 72.3390400151472 | 5.28551875731876 | 59.62055031 | 10419811840275.082 | aggregate | 0.472418297 | 28.16583887 |
| GAK15 | 128 | A3 | 72.3390400151472 | 5.28551875731876 | 59.62055031 | 10419811840275.082 | dense_detritus | 0.023513027 | 1.401859619 |
| GAK15 | 128 | A3 | 72.3390400151472 | 5.28551875731876 | 59.62055031 | 10419811840275.082 | unidentified | 0.048299248 | 2.879627719 |
| GAK15 | 128 | A3 | 72.3390400151472 | 5.28551875731876 | 59.62055031 | 10419811840275.082 | large_loose_pellet | 0.124796973 | 7.440464209 |
| GAK15 | 128 | A3 | 72.3390400151472 | 5.28551875731876 | 59.62055031 | 10419811840275.082 | long_fecal_pellet | 0.081238995 | 4.843513584 |
| GAK15 | 128 | A3 | 72.3390400151472 | 5.28551875731876 | 59.62055031 | 10419811840275.082 | mini_pellet | 0.000965424 | 0.05755909 |
| GAK15 | 128 | A3 | 72.3390400151472 | 5.28551875731876 | 59.62055031 | 10419811840275.082 | salp_pellet | 0.196820598 | 11.73455235 |
| GAK15 | 128 | A3 | 72.3390400151472 | 5.28551875731876 | 59.62055031 | 10419811840275.082 | short_pellet | 0.05107022 | 3.044834636 |
| GAK15 | 128 | A3 | 72.3390400151472 | 5.28551875731876 | 59.62055031 | 10419811840275.082 | phytoplankton | 0 | 0 |
| GAK15 | 128 | A3 | 72.3390400151472 | 5.28551875731876 | 59.62055031 | 10419811840275.082 | rhizaria | 0.000877218 | 0.052300226 |
| GAK15 | 27 | E1 | 232.178291670058 | 76.7226796095952 | 187.7650582 | 72381812861571.16 | aggregate | 0.652757353 | 122.5650225 |
| GAK15 | 27 | E1 | 232.178291670058 | 76.7226796095952 | 187.7650582 | 72381812861571.16 | dense_detritus | 0.018880296 | 3.545059895 |
| GAK15 | 27 | E1 | 232.178291670058 | 76.7226796095952 | 187.7650582 | 72381812861571.16 | unidentified | 0.088396258 | 16.59772845 |
| GAK15 | 27 | E1 | 232.178291670058 | 76.7226796095952 | 187.7650582 | 72381812861571.16 | large_loose_pellet | 0.054196533 | 10.17621511 |

| | | | | | | | | | |
|---|---|---|---|---|---|---|---|---|---|
| GAK15 | 27 | E1 | 232.178291670058 | 76.7226796095952 | 187.7650582 | 72381812861571.16 | long_fecal_pellet | 0.05837099 | 10.96003228 |
| GAK15 | 27 | E1 | 232.178291670058 | 76.7226796095952 | 187.7650582 | 72381812861571.16 | mini_pellet | 0.007175201 | 1.347251971 |
| GAK15 | 27 | E1 | 232.178291670058 | 76.7226796095952 | 187.7650582 | 72381812861571.16 | salp_pellet | 0 | 0 |
| GAK15 | 27 | E1 | 232.178291670058 | 76.7226796095952 | 187.7650582 | 72381812861571.16 | short_pellet | 0.117913302 | 22.13999808 |
| GAK15 | 27 | E1 | 232.178291670058 | 76.7226796095952 | 187.7650582 | 72381812861571.16 | phytoplankton | 0.001757318 | 0.329962884 |
| GAK15 | 27 | E1 | 232.178291670058 | 76.7226796095952 | 187.7650582 | 72381812861571.16 | rhizaria | 0.00055275 | 0.103787101 |
| GAK15 | 27 | E3 | 232.178291670058 | 76.7226796095952 | 167.3545939 | 34409756486962.64 | aggregate | 0.52041271 | 87.09345784 |
| GAK15 | 27 | E3 | 232.178291670058 | 76.7226796095952 | 167.3545939 | 34409756486962.64 | dense_detritus | 0.029253317 | 4.895676949 |
| GAK15 | 27 | E3 | 232.178291670058 | 76.7226796095952 | 167.3545939 | 34409756486962.64 | unidentified | 0.098630367 | 16.50624503 |
| GAK15 | 27 | E3 | 232.178291670058 | 76.7226796095952 | 167.3545939 | 34409756486962.64 | large_loose_pellet | 0.073223801 | 12.25433951 |
| GAK15 | 27 | E3 | 232.178291670058 | 76.7226796095952 | 167.3545939 | 34409756486962.64 | long_fecal_pellet | 0.068536003 | 11.469815 |
| GAK15 | 27 | E3 | 232.178291670058 | 76.7226796095952 | 167.3545939 | 34409756486962.64 | mini_pellet | 0.023271545 | 3.894599888 |
| GAK15 | 27 | E3 | 232.178291670058 | 76.7226796095952 | 167.3545939 | 34409756486962.64 | salp_pellet | 0 | 0 |
| GAK15 | 27 | E3 | 232.178291670058 | 76.7226796095952 | 167.3545939 | 34409756486962.64 | short_pellet | 0.18643301 | 31.20042077 |
| GAK15 | 27 | E3 | 232.178291670058 | 76.7226796095952 | 167.3545939 | 34409756486962.64 | phytoplankton | 0 | 0 |
| GAK15 | 27 | E3 | 232.178291670058 | 76.7226796095952 | 167.3545939 | 34409756486962.64 | rhizaria | 0.000239246 | 0.040038949 |
| GAK15 | 53 | D1 | 123.618645995111 | 25.4452800822746 | 136.7494217 | 28419028017823.03 | aggregate | 0.475369023 | 65.00643897 |
| GAK15 | 53 | D1 | 123.618645995111 | 25.4452800822746 | 136.7494217 | 28419028017823.03 | dense_detritus | 0.013375703 | 1.829119654 |
| GAK15 | 53 | D1 | 123.618645995111 | 25.4452800822746 | 136.7494217 | 28419028017823.03 | unidentified | 0.065163283 | 8.911041287 |
| GAK15 | 53 | D1 | 123.618645995111 | 25.4452800822746 | 136.7494217 | 28419028017823.03 | large_loose_pellet | 0.118016414 | 16.13867636 |
| GAK15 | 53 | D1 | 123.618645995111 | 25.4452800822746 | 136.7494217 | 28419028017823.03 | long_fecal_pellet | 0.158654598 | 21.69592454 |
| GAK15 | 53 | D1 | 123.618645995111 | 25.4452800822746 | 136.7494217 | 28419028017823.03 | mini_pellet | 0.004585238 | 0.627028705 |
| GAK15 | 53 | D1 | 123.618645995111 | 25.4452800822746 | 136.7494217 | 28419028017823.03 | salp_pellet | 0.0437134 | 5.977782217 |
| GAK15 | 53 | D1 | 123.618645995111 | 25.4452800822746 | 136.7494217 | 28419028017823.03 | short_pellet | 0.116698822 | 15.95849642 |
| GAK15 | 53 | D1 | 123.618645995111 | 25.4452800822746 | 136.7494217 | 28419028017823.03 | phytoplankton | 0.004319367 | 0.590670999 |
| GAK15 | 53 | D1 | 123.618645995111 | 25.4452800822746 | 136.7494217 | 28419028017823.03 | rhizaria | 0.000104151 | 0.014242545 |
| GAK15 | 53 | D3 | 123.618645995111 | 25.4452800822746 | 194.5359911 | 48623197508023.32 | aggregate | 0.553750975 | 107.7244947 |
| GAK15 | 53 | D3 | 123.618645995111 | 25.4452800822746 | 194.5359911 | 48623197508023.32 | dense_detritus | 0.027266283 | 5.304273357 |
| GAK15 | 53 | D3 | 123.618645995111 | 25.4452800822746 | 194.5359911 | 48623197508023.32 | unidentified | 0.087635535 | 17.04826557 |
| GAK15 | 53 | D3 | 123.618645995111 | 25.4452800822746 | 194.5359911 | 48623197508023.32 | large_loose_pellet | 0.064842879 | 12.61427366 |
| GAK15 | 53 | D3 | 123.618645995111 | 25.4452800822746 | 194.5359911 | 48623197508023.32 | long_fecal_pellet | 0.148636582 | 28.9151647 |

| | | | | | | | | | |
|---|---|---|---|---|---|---|---|---|---|
| GAK15 | 53 | D3 | 123.618645995111 | 25.4452800822746 | 194.5359911 | 48623197508023.32 | mini_pellet | 0.009483382 | 1.844859072 |
| GAK15 | 53 | D3 | 123.618645995111 | 25.4452800822746 | 194.5359911 | 48623197508023.32 | salp_pellet | 0 | 0 |
| GAK15 | 53 | D3 | 123.618645995111 | 25.4452800822746 | 194.5359911 | 48623197508023.32 | short_pellet | 0.108384366 | 21.08466005 |
| GAK15 | 53 | D3 | 123.618645995111 | 25.4452800822746 | 194.5359911 | 48623197508023.32 | phytoplankton | 0 | 0 |
| GAK15 | 53 | D3 | 123.618645995111 | 25.4452800822746 | 194.5359911 | 48623197508023.32 | rhizaria | 0 | 0 |
| GAK15 | 78 | C1 | 128.659969306109 | 75.3585140318845 | 152.040826 | 40264718014705.56 | aggregate | 0.571930316 | 86.9567576 |
| GAK15 | 78 | C1 | 128.659969306109 | 75.3585140318845 | 152.040826 | 40264718014705.56 | dense_detritus | 0.015508243 | 2.357886036 |
| GAK15 | 78 | C1 | 128.659969306109 | 75.3585140318845 | 152.040826 | 40264718014705.56 | unidentified | 0.142936105 | 21.73212353 |
| GAK15 | 78 | C1 | 128.659969306109 | 75.3585140318845 | 152.040826 | 40264718014705.56 | large_loose_pellet | 0.020670688 | 3.142788442 |
| GAK15 | 78 | C1 | 128.659969306109 | 75.3585140318845 | 152.040826 | 40264718014705.56 | long_fecal_pellet | 0.169898465 | 25.83150297 |
| GAK15 | 78 | C1 | 128.659969306109 | 75.3585140318845 | 152.040826 | 40264718014705.56 | mini_pellet | 0.01616464 | 2.457685287 |
| GAK15 | 78 | C1 | 128.659969306109 | 75.3585140318845 | 152.040826 | 40264718014705.56 | salp_pellet | 0 | 0 |
| GAK15 | 78 | C1 | 128.659969306109 | 75.3585140318845 | 152.040826 | 40264718014705.56 | short_pellet | 0.059886156 | 9.105140601 |
| GAK15 | 78 | C1 | 128.659969306109 | 75.3585140318845 | 152.040826 | 40264718014705.56 | phytoplankton | 0.002956829 | 0.44955877 |
| GAK15 | 78 | C1 | 128.659969306109 | 75.3585140318845 | 152.040826 | 40264718014705.56 | rhizaria | 4.86E-05 | 0.007382787 |
| GAK15 | 78 | C3 | 128.659969306109 | 75.3585140318845 | 93.53148382 | 19774824323886.906 | aggregate | 0.46644383 | 43.62718353 |
| GAK15 | 78 | C3 | 128.659969306109 | 75.3585140318845 | 93.53148382 | 19774824323886.906 | dense_detritus | 0.018591386 | 1.73887993 |
| GAK15 | 78 | C3 | 128.659969306109 | 75.3585140318845 | 93.53148382 | 19774824323886.906 | unidentified | 0.188809775 | 17.65965839 |
| GAK15 | 78 | C3 | 128.659969306109 | 75.3585140318845 | 93.53148382 | 19774824323886.906 | large_loose_pellet | 0.182475878 | 17.06723963 |
| GAK15 | 78 | C3 | 128.659969306109 | 75.3585140318845 | 93.53148382 | 19774824323886.906 | long_fecal_pellet | 0.062475138 | 5.843392344 |
| GAK15 | 78 | C3 | 128.659969306109 | 75.3585140318845 | 93.53148382 | 19774824323886.906 | mini_pellet | 0.008446662 | 0.790028826 |
| GAK15 | 78 | C3 | 128.659969306109 | 75.3585140318845 | 93.53148382 | 19774824323886.906 | salp_pellet | 0.003877988 | 0.362713963 |
| GAK15 | 78 | C3 | 128.659969306109 | 75.3585140318845 | 93.53148382 | 19774824323886.906 | short_pellet | 0.051997775 | 4.863429065 |
| GAK15 | 78 | C3 | 128.659969306109 | 75.3585140318845 | 93.53148382 | 19774824323886.906 | phytoplankton | 0.000838945 | 0.078467809 |
| GAK15 | 78 | C3 | 128.659969306109 | 75.3585140318845 | 93.53148382 | 19774824323886.906 | rhizaria | 0.016042623 | 1.500490332 |
| GAK5 | 103 | A1 | 206.435432000362 | 41.5313795878675 | 183.1681017 | 27730393905830.734 | aggregate | 0.698787242 | 127.9955326 |
| GAK5 | 103 | A1 | 206.435432000362 | 41.5313795878675 | 183.1681017 | 27730393905830.734 | dense_detritus | 0.007841459 | 1.436305176 |
| GAK5 | 103 | A1 | 206.435432000362 | 41.5313795878675 | 183.1681017 | 27730393905830.734 | unidentified | 0.071443914 | 13.08624607 |
| GAK5 | 103 | A1 | 206.435432000362 | 41.5313795878675 | 183.1681017 | 27730393905830.734 | large_loose_pellet | 0.075516442 | 13.83220339 |
| GAK5 | 103 | A1 | 206.435432000362 | 41.5313795878675 | 183.1681017 | 27730393905830.734 | long_fecal_pellet | 0.083683423 | 15.32813378 |
| GAK5 | 103 | A1 | 206.435432000362 | 41.5313795878675 | 183.1681017 | 27730393905830.734 | mini_pellet | 0.006245992 | 1.144066552 |

| | | | | | | | | | |
|---|---|---|---|---|---|---|---|---|---|
| GAK5 | 103 | A1 | 206.435432000362 | 41.5313795878675 | 183.1681017 | 27730393905830.734 | salp_pellet | 0.018513445 | 3.391072597 |
| GAK5 | 103 | A1 | 206.435432000362 | 41.5313795878675 | 183.1681017 | 27730393905830.734 | short_pellet | 0.030283492 | 5.546969723 |
| GAK5 | 103 | A1 | 206.435432000362 | 41.5313795878675 | 183.1681017 | 27730393905830.734 | phytoplankton | 0.006224162 | 1.140068012 |
| GAK5 | 103 | A1 | 206.435432000362 | 41.5313795878675 | 183.1681017 | 27730393905830.734 | rhizaria | 0.001460428 | 0.267503767 |
| GAK5 | 27 | D1 | 359.570521068969 | 36.8013171804428 | 101.3579664 | 13609570639367.229 | aggregate | 0.557912236 | 56.54884966 |
| GAK5 | 27 | D1 | 359.570521068969 | 36.8013171804428 | 101.3579664 | 13609570639367.229 | dense_detritus | 0.059189044 | 5.999281156 |
| GAK5 | 27 | D1 | 359.570521068969 | 36.8013171804428 | 101.3579664 | 13609570639367.229 | unidentified | 0.129575099 | 13.13346849 |
| GAK5 | 27 | D1 | 359.570521068969 | 36.8013171804428 | 101.3579664 | 13609570639367.229 | large_loose_pellet | 0.02521756 | 2.556000612 |
| GAK5 | 27 | D1 | 359.570521068969 | 36.8013171804428 | 101.3579664 | 13609570639367.229 | long_fecal_pellet | 0.195042366 | 19.76909762 |
| GAK5 | 27 | D1 | 359.570521068969 | 36.8013171804428 | 101.3579664 | 13609570639367.229 | mini_pellet | 0.014891429 | 1.509364965 |
| GAK5 | 27 | D1 | 359.570521068969 | 36.8013171804428 | 101.3579664 | 13609570639367.229 | salp_pellet | 0 | 0 |
| GAK5 | 27 | D1 | 359.570521068969 | 36.8013171804428 | 101.3579664 | 13609570639367.229 | short_pellet | 0.012526628 | 1.26967358 |
| GAK5 | 27 | D1 | 359.570521068969 | 36.8013171804428 | 101.3579664 | 13609570639367.229 | phytoplankton | 0.0042258 | 0.428318461 |
| GAK5 | 27 | D1 | 359.570521068969 | 36.8013171804428 | 101.3579664 | 13609570639367.229 | rhizaria | 0.001419837 | 0.143911829 |
| GAK5 | 53 | C1 | 160.31688729156 | 17.3907862529961 | 161.5808806 | 22762532901051.375 | aggregate | 0.552664197 | 89.29996763 |
| GAK5 | 53 | C1 | 160.31688729156 | 17.3907862529961 | 161.5808806 | 22762532901051.375 | dense_detritus | 0.072399161 | 11.69832015 |
| GAK5 | 53 | C1 | 160.31688729156 | 17.3907862529961 | 161.5808806 | 22762532901051.375 | unidentified | 0.109694527 | 17.72453831 |
| GAK5 | 53 | C1 | 160.31688729156 | 17.3907862529961 | 161.5808806 | 22762532901051.375 | large_loose_pellet | 0.043346292 | 7.003932106 |
| GAK5 | 53 | C1 | 160.31688729156 | 17.3907862529961 | 161.5808806 | 22762532901051.375 | long_fecal_pellet | 0.159171469 | 25.71906615 |
| GAK5 | 53 | C1 | 160.31688729156 | 17.3907862529961 | 161.5808806 | 22762532901051.375 | mini_pellet | 0.024992628 | 4.038330802 |
| GAK5 | 53 | C1 | 160.31688729156 | 17.3907862529961 | 161.5808806 | 22762532901051.375 | salp_pellet | 0 | 0 |
| GAK5 | 53 | C1 | 160.31688729156 | 17.3907862529961 | 161.5808806 | 22762532901051.375 | short_pellet | 0.034442968 | 5.565325156 |
| GAK5 | 53 | C1 | 160.31688729156 | 17.3907862529961 | 161.5808806 | 22762532901051.375 | phytoplankton | 0.003088429 | 0.499031121 |
| GAK5 | 53 | C1 | 160.31688729156 | 17.3907862529961 | 161.5808806 | 22762532901051.375 | rhizaria | 0.000200328 | 0.032369166 |
| GAK5 | 78 | B1 | 185.873317408334 | 65.6850059588525 | 127.2447589 | 19102493634079.492 | aggregate | 0.515612641 | 65.60900624 |
| GAK5 | 78 | B1 | 185.873317408334 | 65.6850059588525 | 127.2447589 | 19102493634079.492 | dense_detritus | 0.03096544 | 3.940189925 |
| GAK5 | 78 | B1 | 185.873317408334 | 65.6850059588525 | 127.2447589 | 19102493634079.492 | unidentified | 0.054778507 | 6.970277977 |
| GAK5 | 78 | B1 | 185.873317408334 | 65.6850059588525 | 127.2447589 | 19102493634079.492 | large_loose_pellet | 0.149346612 | 19.00357364 |
| GAK5 | 78 | B1 | 185.873317408334 | 65.6850059588525 | 127.2447589 | 19102493634079.492 | long_fecal_pellet | 0.218329854 | 27.78132962 |
| GAK5 | 78 | B1 | 185.873317408334 | 65.6850059588525 | 127.2447589 | 19102493634079.492 | mini_pellet | 0.00133166 | 0.169446796 |
| GAK5 | 78 | B1 | 185.873317408334 | 65.6850059588525 | 127.2447589 | 19102493634079.492 | salp_pellet | 0.011711662 | 1.490247642 |

| | | | | | | | | | |
|---|---|---|---|---|---|---|---|---|---|
| GAK5 | 78 | B1 | 185.873317408334 | 65.6850059588525 | 127.2447589 | 19102493634079.492 | short_pellet | 0.005437709 | 0.69191995 |
| GAK5 | 78 | B1 | 185.873317408334 | 65.6850059588525 | 127.2447589 | 19102493634079.492 | phytoplankton | 0.008321403 | 1.058854957 |
| GAK5 | 78 | B1 | 185.873317408334 | 65.6850059588525 | 127.2447589 | 19102493634079.492 | rhizaria | 0.004164511 | 0.529912185 |
| GAK9 | 103 | B3 | 98.0264136823931 | 10.2895781275852 | 254.870107 | 53737125822853.3 | aggregate | 0.770907295 | 196.4812247 |
| GAK9 | 103 | B3 | 98.0264136823931 | 10.2895781275852 | 254.870107 | 53737125822853.3 | dense_detritus | 0.012089328 | 3.081208251 |
| GAK9 | 103 | B3 | 98.0264136823931 | 10.2895781275852 | 254.870107 | 53737125822853.3 | unidentified | 0.050736707 | 12.93126988 |
| GAK9 | 103 | B3 | 98.0264136823931 | 10.2895781275852 | 254.870107 | 53737125822853.3 | large_loose_pellet | 0.036053275 | 9.18890203 |
| GAK9 | 103 | B3 | 98.0264136823931 | 10.2895781275852 | 254.870107 | 53737125822853.3 | long_fecal_pellet | 0.07989651 | 20.36323198 |
| GAK9 | 103 | B3 | 98.0264136823931 | 10.2895781275852 | 254.870107 | 53737125822853.3 | mini_pellet | 0.009922124 | 2.528852725 |
| GAK9 | 103 | B3 | 98.0264136823931 | 10.2895781275852 | 254.870107 | 53737125822853.3 | salp_pellet | 0 | 0 |
| GAK9 | 103 | B3 | 98.0264136823931 | 10.2895781275852 | 254.870107 | 53737125822853.3 | short_pellet | 0.03913353 | 9.973967093 |
| GAK9 | 103 | B3 | 98.0264136823931 | 10.2895781275852 | 254.870107 | 53737125822853.3 | phytoplankton | 0.001245065 | 0.317329939 |
| GAK9 | 103 | B3 | 98.0264136823931 | 10.2895781275852 | 254.870107 | 53737125822853.3 | rhizaria | 1.62E-05 | 0.004120397 |
| GAK9 | 128 | A1 | 85.8744866751674 | 8.97054169464314 | 194.6077885 | 34744489466554.9 | aggregate | 0.672440997 | 130.8622554 |
| GAK9 | 128 | A1 | 85.8744866751674 | 8.97054169464314 | 194.6077885 | 34744489466554.9 | dense_detritus | 0.136375325 | 26.53970035 |
| GAK9 | 128 | A1 | 85.8744866751674 | 8.97054169464314 | 194.6077885 | 34744489466554.9 | unidentified | 0.028544444 | 5.55497116 |
| GAK9 | 128 | A1 | 85.8744866751674 | 8.97054169464314 | 194.6077885 | 34744489466554.9 | large_loose_pellet | 0.050830574 | 9.892025558 |
| GAK9 | 128 | A1 | 85.8744866751674 | 8.97054169464314 | 194.6077885 | 34744489466554.9 | long_fecal_pellet | 0.087709136 | 17.06888105 |
| GAK9 | 128 | A1 | 85.8744866751674 | 8.97054169464314 | 194.6077885 | 34744489466554.9 | mini_pellet | 0.007090005 | 1.379770194 |
| GAK9 | 128 | A1 | 85.8744866751674 | 8.97054169464314 | 194.6077885 | 34744489466554.9 | salp_pellet | 0.002009748 | 0.391112676 |
| GAK9 | 128 | A1 | 85.8744866751674 | 8.97054169464314 | 194.6077885 | 34744489466554.9 | short_pellet | 0.01318616 | 2.56612946 |
| GAK9 | 128 | A1 | 85.8744866751674 | 8.97054169464314 | 194.6077885 | 34744489466554.9 | phytoplankton | 0.00146392 | 0.284890157 |
| GAK9 | 128 | A1 | 85.8744866751674 | 8.97054169464314 | 194.6077885 | 34744489466554.9 | rhizaria | 0.000349691 | 0.068052518 |
| GAK9 | 27 | E1 | 320.233405920397 | 122.334639077153 | 116.2659074 | 19756718396091.17 | aggregate | 0.712219901 | 82.80689304 |
| GAK9 | 27 | E1 | 320.233405920397 | 122.334639077153 | 116.2659074 | 19756718396091.17 | dense_detritus | 0.017929619 | 2.084603433 |
| GAK9 | 27 | E1 | 320.233405920397 | 122.334639077153 | 116.2659074 | 19756718396091.17 | unidentified | 0.098039368 | 11.39863604 |
| GAK9 | 27 | E1 | 320.233405920397 | 122.334639077153 | 116.2659074 | 19756718396091.17 | large_loose_pellet | 0.008147974 | 0.947331618 |
| GAK9 | 27 | E1 | 320.233405920397 | 122.334639077153 | 116.2659074 | 19756718396091.17 | long_fecal_pellet | 0.104085799 | 12.10162984 |
| GAK9 | 27 | E1 | 320.233405920397 | 122.334639077153 | 116.2659074 | 19756718396091.17 | mini_pellet | 0.023665114 | 2.751445984 |
| GAK9 | 27 | E1 | 320.233405920397 | 122.334639077153 | 116.2659074 | 19756718396091.17 | salp_pellet | 0 | 0 |
| GAK9 | 27 | E1 | 320.233405920397 | 122.334639077153 | 116.2659074 | 19756718396091.17 | short_pellet | 0.030805747 | 3.581658139 |

| | | | | | | | | | |
|---|---|---|---|---|---|---|---|---|---|
| GAK9 | 27 | E1 | 320.233405920397 | 122.334639077153 | 116.2659074 | 19756718396091.17 | phytoplankton | 0.003217986 | 0.374142047 |
| GAK9 | 27 | E1 | 320.233405920397 | 122.334639077153 | 116.2659074 | 19756718396091.17 | rhizaria | 0.001888492 | 0.219567266 |
| GAK9 | 53 | D1 | 143.881147588412 | 35.0525194601771 | 49.89177913 | 8193633587491.709 | aggregate | 0.517027801 | 25.79543686 |
| GAK9 | 53 | D1 | 143.881147588412 | 35.0525194601771 | 49.89177913 | 8193633587491.709 | dense_detritus | 0.025856747 | 1.290039129 |
| GAK9 | 53 | D1 | 143.881147588412 | 35.0525194601771 | 49.89177913 | 8193633587491.709 | unidentified | 0.065947927 | 3.290259412 |
| GAK9 | 53 | D1 | 143.881147588412 | 35.0525194601771 | 49.89177913 | 8193633587491.709 | large_loose_pellet | 0.028131782 | 1.403544655 |
| GAK9 | 53 | D1 | 143.881147588412 | 35.0525194601771 | 49.89177913 | 8193633587491.709 | long_fecal_pellet | 0.177630677 | 8.862310523 |
| GAK9 | 53 | D1 | 143.881147588412 | 35.0525194601771 | 49.89177913 | 8193633587491.709 | mini_pellet | 0.071379783 | 3.561264375 |
| GAK9 | 53 | D1 | 143.881147588412 | 35.0525194601771 | 49.89177913 | 8193633587491.709 | salp_pellet | 0 | 0 |
| GAK9 | 53 | D1 | 143.881147588412 | 35.0525194601771 | 49.89177913 | 8193633587491.709 | short_pellet | 0.080056558 | 3.994164133 |
| GAK9 | 53 | D1 | 143.881147588412 | 35.0525194601771 | 49.89177913 | 8193633587491.709 | phytoplankton | 0.006951696 | 0.346832497 |
| GAK9 | 53 | D1 | 143.881147588412 | 35.0525194601771 | 49.89177913 | 8193633587491.709 | rhizaria | 0.027017027 | 1.34792755 |
| GAK9 | 78 | C1 | 114.44658681547 | 10.8403488302991 | 109.5785742 | 19125161177551.406 | aggregate | 0.691405385 | 75.76321632 |
| GAK9 | 78 | C1 | 114.44658681547 | 10.8403488302991 | 109.5785742 | 19125161177551.406 | dense_detritus | 0.022162086 | 2.428489744 |
| GAK9 | 78 | C1 | 114.44658681547 | 10.8403488302991 | 109.5785742 | 19125161177551.406 | unidentified | 0.095637884 | 10.47986298 |
| GAK9 | 78 | C1 | 114.44658681547 | 10.8403488302991 | 109.5785742 | 19125161177551.406 | large_loose_pellet | 0.036761596 | 4.028283279 |
| GAK9 | 78 | C1 | 114.44658681547 | 10.8403488302991 | 109.5785742 | 19125161177551.406 | long_fecal_pellet | 0.117896793 | 12.91896244 |
| GAK9 | 78 | C1 | 114.44658681547 | 10.8403488302991 | 109.5785742 | 19125161177551.406 | mini_pellet | 0.006105781 | 0.669062802 |
| GAK9 | 78 | C1 | 114.44658681547 | 10.8403488302991 | 109.5785742 | 19125161177551.406 | salp_pellet | 0.017946493 | 1.966551115 |
| GAK9 | 78 | C1 | 114.44658681547 | 10.8403488302991 | 109.5785742 | 19125161177551.406 | short_pellet | 0.010582816 | 1.159649844 |
| GAK9 | 78 | C1 | 114.44658681547 | 10.8403488302991 | 109.5785742 | 19125161177551.406 | phytoplankton | 0.001239656 | 0.135839784 |
| GAK9 | 78 | C1 | 114.44658681547 | 10.8403488302991 | 109.5785742 | 19125161177551.406 | rhizaria | 0.00026151 | 0.028655881 |
| KOD10 | 53 | D3 | 481.959933707405 | 138.497339467903 | 147.6941818 | 37576086681139.04 | aggregate | 0.808074284 | 119.3478703 |
| KOD10 | 53 | D3 | 481.959933707405 | 138.497339467903 | 147.6941818 | 37576086681139.04 | dense_detritus | 0.035313168 | 5.215549427 |
| KOD10 | 53 | D3 | 481.959933707405 | 138.497339467903 | 147.6941818 | 37576086681139.04 | unidentified | 0.025477677 | 3.762904597 |
| KOD10 | 53 | D3 | 481.959933707405 | 138.497339467903 | 147.6941818 | 37576086681139.04 | large_loose_pellet | 0.020367721 | 3.008193926 |
| KOD10 | 53 | D3 | 481.959933707405 | 138.497339467903 | 147.6941818 | 37576086681139.04 | long_fecal_pellet | 0.070281035 | 10.38010002 |
| KOD10 | 53 | D3 | 481.959933707405 | 138.497339467903 | 147.6941818 | 37576086681139.04 | mini_pellet | 0.001726117 | 0.254937455 |
| KOD10 | 53 | D3 | 481.959933707405 | 138.497339467903 | 147.6941818 | 37576086681139.04 | salp_pellet | 0.002817143 | 0.41607564 |
| KOD10 | 53 | D3 | 481.959933707405 | 138.497339467903 | 147.6941818 | 37576086681139.04 | short_pellet | 0.034411865 | 5.082432188 |
| KOD10 | 53 | D3 | 481.959933707405 | 138.497339467903 | 147.6941818 | 37576086681139.04 | phytoplankton | 0.00153099 | 0.226118316 |

| | | | | | | | | | |
|---|---|---|---|---|---|---|---|---|---|
| KOD10 | 53 | D3 | 481.959933707405 | 138.497339467903 | 147.6941818 | 37576086681139.04 | rhizaria | 0 | 0 |
| KOD5 | 27 | B3 | 1038.54327352925 | 82.8246386828398 | 753.9042174 | 260480053311654.66 | aggregate | 0.933303227 | 703.6212386 |
| KOD5 | 27 | B3 | 1038.54327352925 | 82.8246386828398 | 753.9042174 | 260480053311654.66 | dense_detritus | 0.005130872 | 3.868185892 |
| KOD5 | 27 | B3 | 1038.54327352925 | 82.8246386828398 | 753.9042174 | 260480053311654.66 | unidentified | 0.021580036 | 16.26927997 |
| KOD5 | 27 | B3 | 1038.54327352925 | 82.8246386828398 | 753.9042174 | 260480053311654.66 | large_loose_pellet | 0.011533291 | 8.694996472 |
| KOD5 | 27 | B3 | 1038.54327352925 | 82.8246386828398 | 753.9042174 | 260480053311654.66 | long_fecal_pellet | 0.006140324 | 4.629216115 |
| KOD5 | 27 | B3 | 1038.54327352925 | 82.8246386828398 | 753.9042174 | 260480053311654.66 | mini_pellet | 0.011033389 | 8.318118225 |
| KOD5 | 27 | B3 | 1038.54327352925 | 82.8246386828398 | 753.9042174 | 260480053311654.66 | salp_pellet | 0 | 0 |
| KOD5 | 27 | B3 | 1038.54327352925 | 82.8246386828398 | 753.9042174 | 260480053311654.66 | short_pellet | 0.009313102 | 7.021187068 |
| KOD5 | 27 | B3 | 1038.54327352925 | 82.8246386828398 | 753.9042174 | 260480053311654.66 | phytoplankton | 0.001722198 | 1.298372559 |
| KOD5 | 27 | B3 | 1038.54327352925 | 82.8246386828398 | 753.9042174 | 260480053311654.66 | rhizaria | 0.000243562 | 0.183622503 |

15

**Supporting Information Figures**

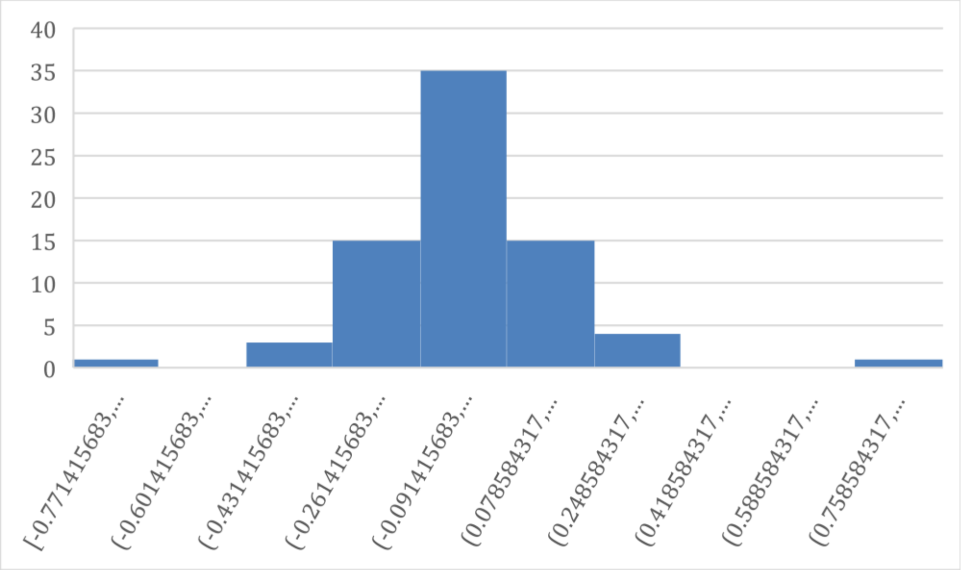

Supporting Information Fig. 1. Relative residuals of particulate organic carbon (POC) collected in paired sediment trap tubes at the same depth.

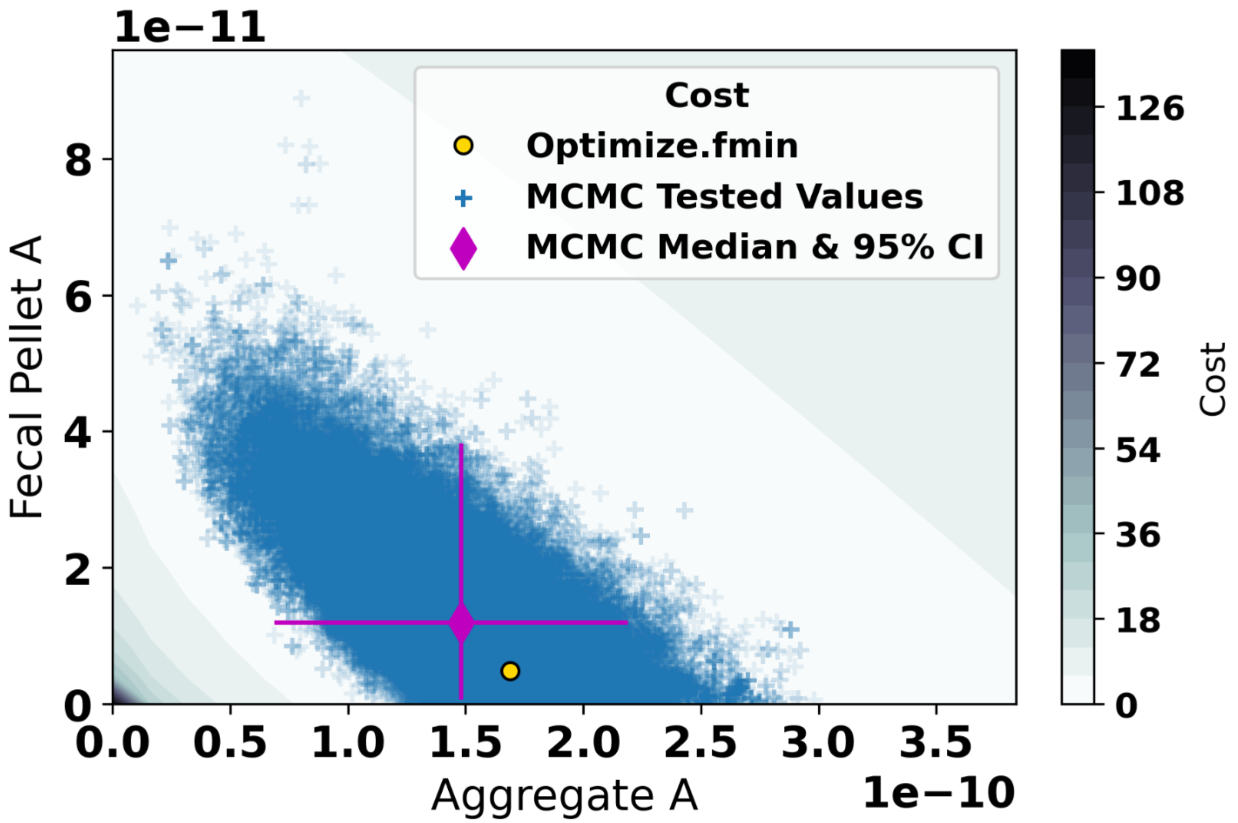

Supporting Information Fig. 2. Constraining Aggregate A and Fecal Pellet A values. Each value tested with the Markov chain Monte Carlo (MCMC) method (blue plus signs) is shown along with the median value (magenta diamond) and the 95% confidence interval. The starting value for the MCMC is the result from Optimize.fmin (yellow circle) is also plotted. The overall cost (residual r squared) space is plotted as contours in the background.

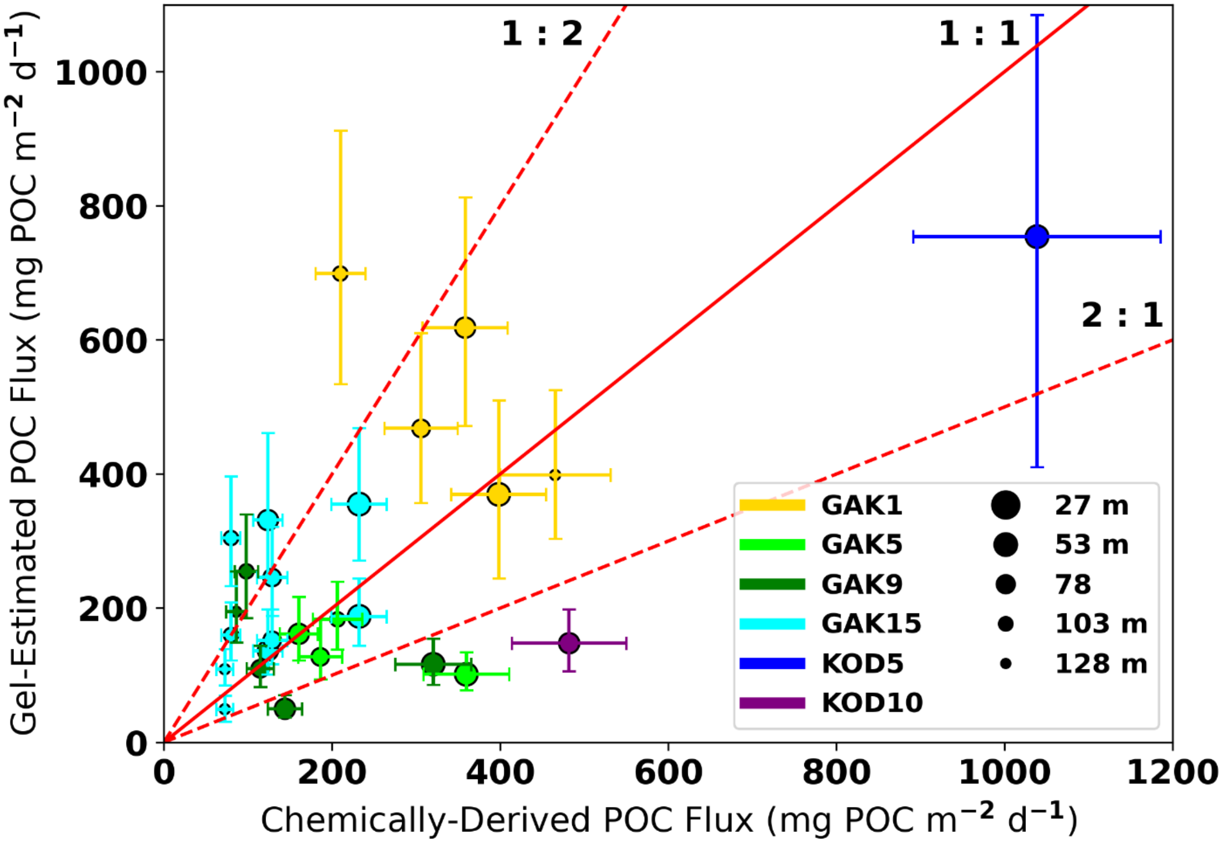

Supporting Information Fig. 3. Chemically-derived formalin-preserved bulk sinking particulate organic carbon (POC) flux versus gel-derived estimated POC flux from particle images. The red line shows the 1:1 relationship and the dashed lines show the 2:1 or 1:2 relationship. Horizontal error bars represent an average chemically-derived POC flux error of approximately 14%. Vertical error bars represent the 95% confidence interval from the Markov chain Monte Carlo analysis.

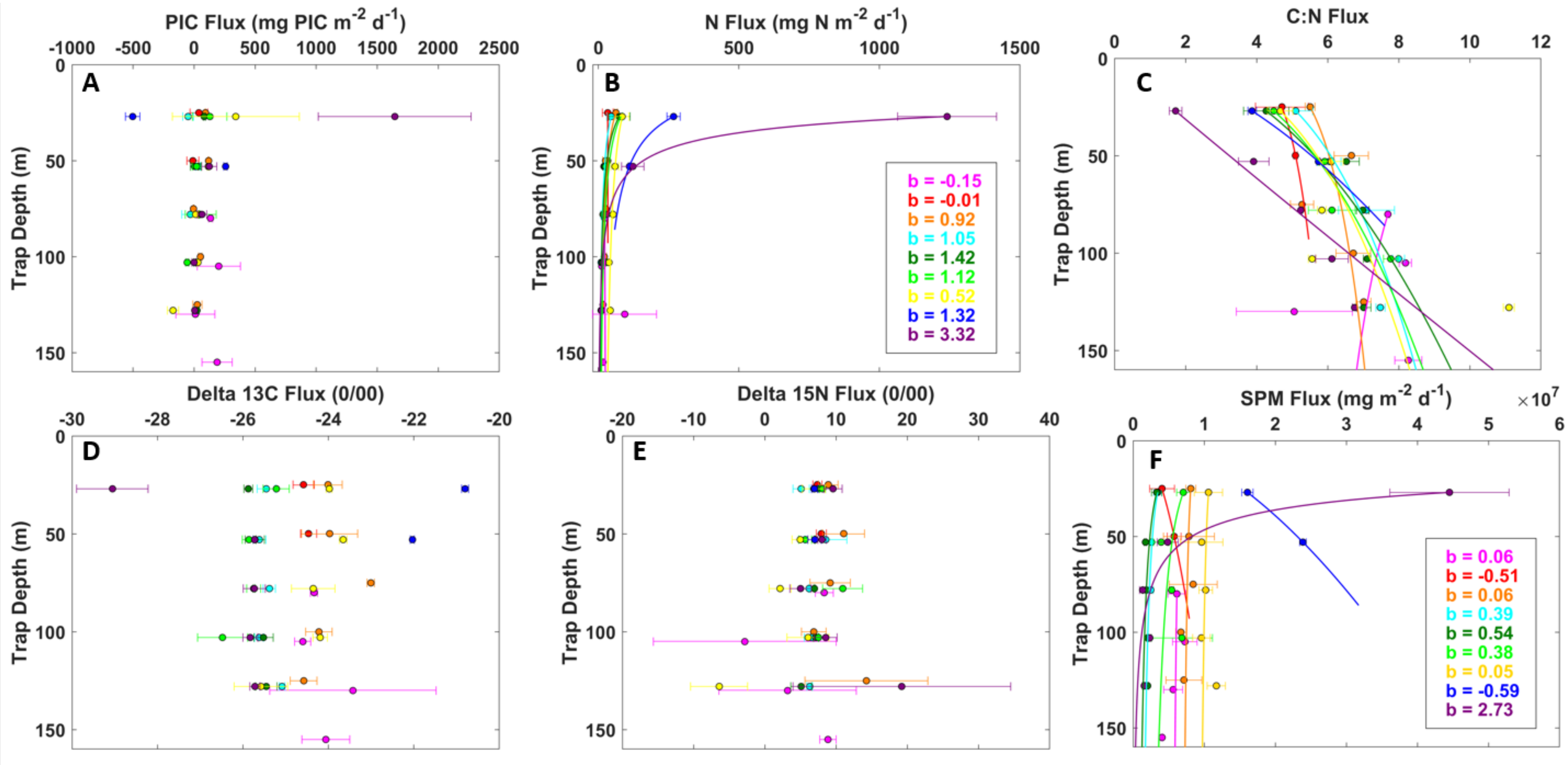

Supporting Information Fig. 4. (A) Particulate inorganic carbon (PIC) flux (mg PIC $m^{-2}$ $day^{-1}$) over depth (m). (B) Particulate nitrogen (N) flux (mg N $m^{-2}$ $day^{-1}$) over depth (m). (C) Carbon to Nitrogen (C:N) ratio over depth (m). (D) Delta 13C flux (0/00) over depth (m). (E) Delta 15N flux (0/00) over depth (m). (F) Total suspended particulate matter (SPM, mg) over depth (m). Delta $^{15}$N and $\delta^{13}$C stable isotope data were obtained using continuous-flow isotope ratio mass spectrometry. Stable isotope ratios were reported in $\delta$

notation as parts per thousand (‰) deviation from the international standards VPDB (carbon) and air (nitrogen). Typically, instrument precision was <0.2 ‰.

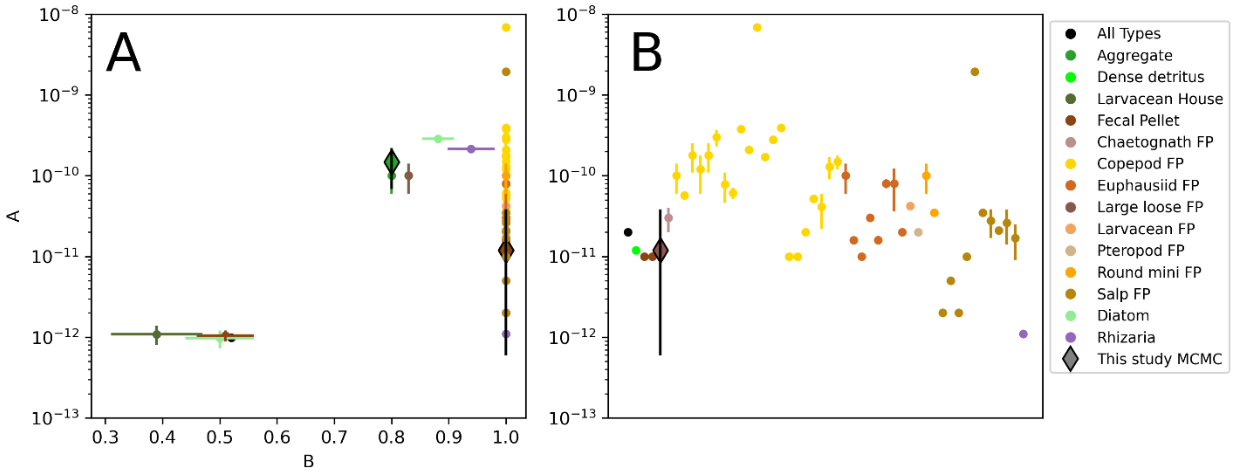

Supporting Information Fig. 5. (A) Empirical and modeled values of A and B values for the equation $carbon = A \times volume^{B}$ taken from various studies (circle-shaped markers, see Supplemental Table 2 for values and references). The results from the gel-derived POC flux from this study (Markov chain Monte Carlo, diamond-shaped markers with 95% confidence interval, black error bar) are within the range of empirical values. (B) All A values where B = 1 are plotted sequentially by type for clarity. FP = fecal pellet. See Supplementary Table 4 for actual data and sources.

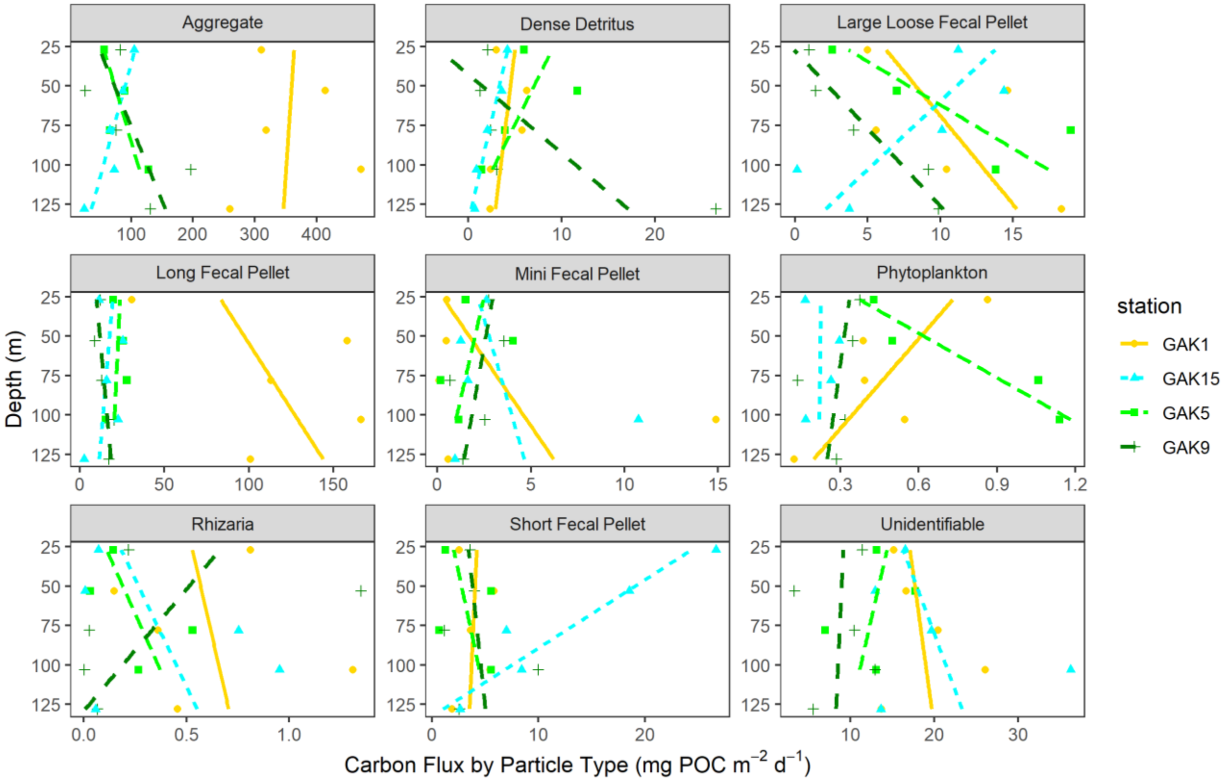

Supporting Information Fig. 6. Gel-estimated carbon flux over depth separated by particle type and colored by station fitted with a linear curve to show changes in gel estimate carbon flux by type over depth.

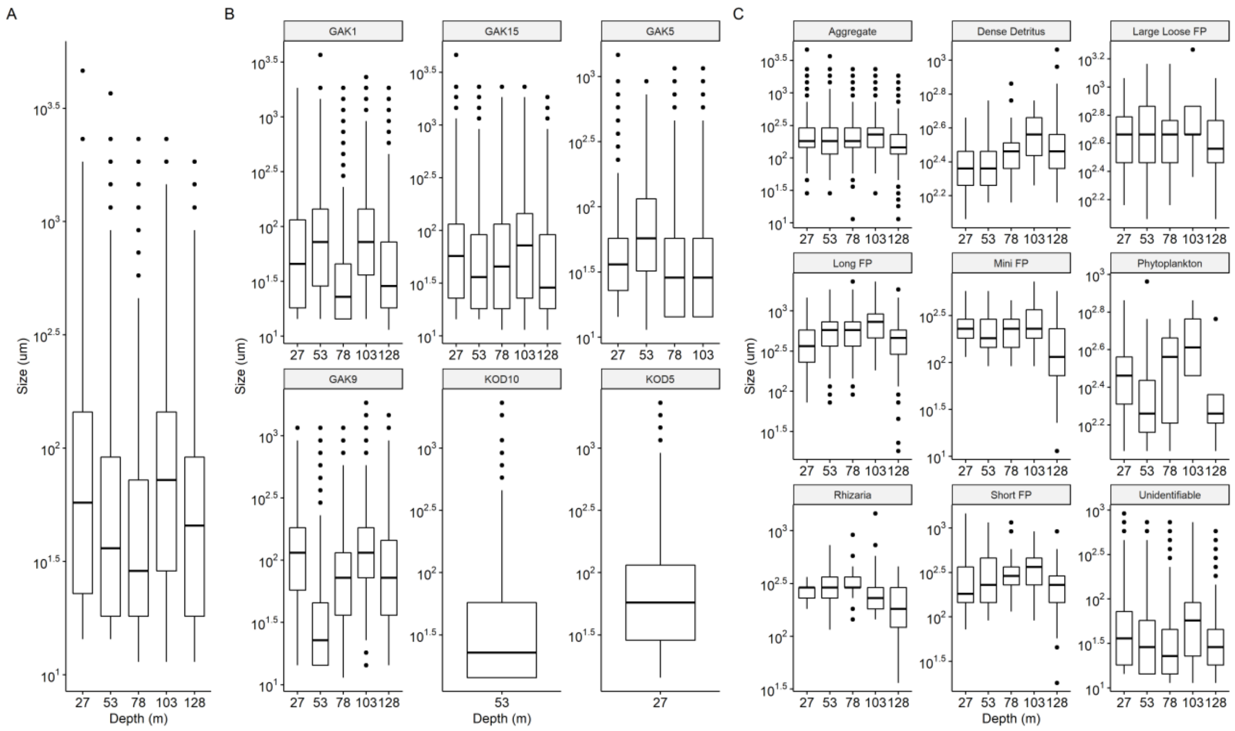

Supporting Information Fig. 7. Box and whisker plots showing mean (line within box) and interquartile range (box) of all sinking particle sizes detected from gel analysis over depth (A) on a log scale with only stations sampled at all 5 sampling depths included, (B) on a log scale separated by station, and (D) on a log scale separated by particle type, with only stations sampled at all 5 sampling depths included.

### Citations and References

Abou Debs, C. 1984. Carbon and nitrogen budget of the calanoid copepod Temora stylifera: effect of concentration and composition of food. Mar Ecol Prog Ser **15**: 213–223.

Alldredge, A. 1998. The carbon, nitrogen and mass content of marine snow as a function of aggregate size. Deep Sea Research I **45**: 529–541.

Belcher, A., M. Iversen, C. Manno, S. A. Henson, G. A. Tarling, and R. Sanders. 2016. The role of particle associated microbes in remineralization of fecal pellets in the upper mesopelagic of the Scotia Sea, Antarctica. Limnol Oceanogr **61**: 1049–1064. doi:10.1002/lno.10269

Butler, M., and H. G. Dam. 1994. Production rates and characteristics of fecal pellets of the copepod Acartia tonsa under simulated phytoplankton bloom conditions: Implications for vertical fluxes. Mar Ecol Prog Ser **114**: 81–92. doi:10.3354/meps114081

California Current Ecosystem LTER, and G. Goericke. 2022. Water column primary production per day integrated over the euphotic zone from CCE LTER process cruises in the California Current System, 2006 - 2019 (ongoing). ver 5. Environmental Data Initiative. doi:https://doi.org/10.6073/pasta/9dddc0e6ab3c0ac48d41e04811a1870f (Accessed 2023-02-28)

California Current Ecosystem LTER, M. Stukel, and M. Landry. 2022. Exported particulate carbon and nitrogen measurements from 4-day sediment trap deployments in the CCE region, 2007 - 2019 (ongoing). ver 7. Environmental Data Initiative.

Caron, D. A., L. P. Madin, and J. J. Cole. 1989. Composition and degradation of salp fecal pellets: implications for vertical flux in oceanic environments. J Mar Res **47**: 829–850. doi:10.1357/002224089785076118

Carroll, M. L., J.-C. Miquel, and S. W. Fowler. 1998. Seasonal patterns and depth-specific trends of zooplankton fecal pellet fluxes in the Northwestern Mediterranean Sea.

Durkin, C. A., K. O. Buesseler, I. Cetinić, M. L. Estapa, R. P. Kelly, and M. Omand. 2021. A Visual Tour of Carbon Export by Sinking Particles. Global Biogeochem Cycles **35**: 1–17. doi:10.1029/2021GB006985

Gleiber, M. R., D. K. Steinberg, and H. W. Ducklow. 2012. Time series of vertical flux of zooplankton fecal pellets on the continental shelf of the western Antarctic Peninsula. Mar Ecol Prog Ser **471**: 23–36. doi:10.3354/meps10021

Gonzalez, H. E. 1992. The distribution and abundance of krill faecal material and oval pellets in the Scotia and Weddell Seas ( Antarctica) and their role in particle flux. Polar Biol **12**: 81–91.

Gonzalez, H. E., S. R. Gonzalez, and G. J. A. Brummer. 1994. Short-term sedimentation pattern of zooplankton, faeces and microplankton at a permanent station in the Bjornafjorden (Norway) during April May 1992. Mar Ecol Prog Ser **105**: 31–46. doi:10.3354/meps105031

Gowing, M. M., D. L. Garrison, H. B. Kunze, and C. J. Winchell. 2001. Biological components of Ross Sea short-term particle fluxes in the austral summer of1995–1996. Deep-Sea Research I **48**: 2645–2671.

Hansen, B., F. L. Fotel, N. J. Jensen, and S. D. Madsen. 1996. Bacteria associated with a marine planktonic copepod in culture. II. Degradation of fecal pellets produced on a diatom, a nanoflagellate or a dinoflagellate diet. J Plankton Res **18**: 275–288. doi:10.1093/plankt/18.2.275

Honjo, S., and M. R. Roman. 1978. Marine copepod fecal pellets: production, preservation and sedimentation. J Mar Res **36**: 45–57.

Huskin, I., L. Viesca, and R. Anadón. 2004. Particle flux in the Subtropical Atlantic near the Azores: Influence of mesozooplankton. J Plankton Res **26**: 403–415. doi:10.1093/plankt/fbh031

Iversen, M. H., E. A. Pakhomov, B. P. V. Hunt, H. van der Jagt, D. Wolf-Gladrow, and C. Klaas. 2017. Sinkers or floaters? Contribution from salp pellets to the export flux during a large bloom event in the Southern Ocean. Deep Sea Res 2 Top Stud Oceanogr **138**: 116–125. doi:10.1016/j.dsr2.2016.12.004

Kahru, M., Z. Lee, R. M. Kudela, M. Manzano-Sarabia, and B. Greg Mitchell. 2015. Multi-satellite time series of inherent optical properties in the California Current. Deep Sea Res 2 Top Stud Oceanogr **112**: 91–106. doi:10.1016/j.dsr2.2013.07.023

Lane, P. V. Z., S. L. Smith, J. L. Urban, and P. E. Biscaye. 1994. Carbon flux and recycling associated with zooplanktonic fecal pellets on the shelf of the Middle Atlantic Bight. Deep Sea Research Part A **41**: 437–457.

Lundsgaard, C., and M. Olesen. 1997. The Origin of Sedimenting Detrital Matter in a Coastal System. Limnol Oceanogr **42**: 1001–1005.

Madin, L. P. 1982. Production, Composition and Sedimentation of Salp Fecal Pellets in Oceanic Waters. Mar Biol **67**: 39–45.

Manno, C., G. Stowasser, P. Enderlein, S. Fielding, and G. A. Tarling. 2015. The contribution of zooplankton faecal pellets to deep-carbon transport in the Scotia Sea (Southern Ocean). Biogeosciences **12**: 1955–1965. doi:10.5194/bg-12-1955-2015

Menden-Deuer, S., and E. J. Lessard. 2000. Carbon to volume relationships for dinoflagellates, diatoms, and other protist plankton. Limnol Oceanogr **45**: 569–579. doi:10.4319/lo.2000.45.3.0569

Olesen, M., S. Strake, and A. Andrushaitis. 2005. Egestion of non-pellet-bound fecal material from the copepod Acartia tonsa: Implication for vertical flux and degradation. Mar Ecol Prog Ser **293**: 131–142. doi:10.3354/meps293131

Pauli, N. C., C. M. Flintrop, C. Konrad, and others. 2021. Krill and salp faecal pellets contribute equally to the carbon flux at the Antarctic Peninsula. Nat Commun **12**. doi:10.1038/s41467-021-27436-9

Reigstad, M., C. Wexels Riser, and C. Svensen. 2005. Fate of copepod faecal pellets and the role of Oithona spp. Mar Ecol Prog Ser **304**: 265–270.

Roy, S., N. Silverberg, N. Romero, and others. 2000. Importance of mesozooplankton feeding for the downward flux of biogenic carbon in the Gulf of St. Lawrence (Canada). Deep-Sea Research II **47**: 519–544.

Silver, M. W., and K. W. Bruland. 1981. Differential feeding and fecal pellet composition of salps and pteropods, and the possible origin of the deep-water flora and olive-green "Cells." Mar Biol **62**: 263–273. doi:10.1007/BF00397693

Silver, M. W., and M. M. Gowing. 1991. The "Particle" Flux: Origins and biological components. Prog Oceanogr **26**: 75–113.

Smith, W. O., A. R. Shields, J. C. Dreyer, J. A. Peloquin, and V. Asper. 2011. Interannual variability in vertical export in the Ross Sea: Magnitude, composition, and environmental correlates. Deep Sea Res 1 Oceanogr Res Pap **58**: 147–159. doi:10.1016/j.dsr.2010.11.007

Stukel, M. R., T. Biard, J. Krause, and M. D. Ohman. 2018. Large Phaeodaria in the twilight zone: Their role in the carbon cycle. Limnol Oceanogr **63**: 2579–2594. doi:10.1002/lno.10961

Suzuki, H., H. Sasaki, and M. Fukuchi. 2003. Loss Processes of Sinking Fecal Pellets of Zooplankton in the Mesopelagic Layers of the Antarctic Marginal Ice Zone. J Oceanogr **59**.

Taguchi, S., and T. Saino. 1998. Net Zooplankton and the Biological Pump off Sanriku, Japan. J Oceanogr **54**: 573–582.

Turner, J. T. 2015. Zooplankton fecal pellets, marine snow, phytodetritus and the ocean's biological pump. Prog Oceanogr **130**: 205–248. doi:10.1016/j.pocean.2014.08.005

Urban-Rich, J. 1997. Latitudinal variations in copepod fecal pellet contribution to organic carbon and amino acid flux. PhD dissertation, University of Maryland, College Park.

Urban-Rich, J., D. A. Hansell, and M. R. Roman. 1998. Analysis of copepod fecal pellet carbon using a high temperature combustion method. Mar Ecol Prog Ser **171**: 199–208. doi:10.3354/meps171199

Urrère, M. A., and G. A. Knauer. 1981. Zooplankton fecal pellet fluxes and vertical transport of particulate organic material in the pelagic environment. J Plankton Res **3**: 369–387. doi:10.1093/plankt/3.3.369

Wassmann, P., J. E. Ypma, and A. Tselepides. 2000. Vertical flux of faecal pellets and microplankton on the shelf of the oligotrophic Cretan Sea (NE Mediterranean Sea). Prog Oceanogr **46**: 241–258.

Wexels Riser, C., P. Wassmann, K. Olli, and E. Arashkevich. 2001. Production, retention and export of zooplankton faecal pellets on and off the Iberian shelf, north-west Spain. Prog Oceanogr **51**: 423–441.

Wexels Riser, C., P. Wassmann, M. Reigstad, and L. Seuthe. 2008. Vertical flux regulation by zooplankton in the northern Barents Sea during Arctic spring. Deep Sea Res 2 Top Stud Oceanogr **55**: 2320–2329. doi:10.1016/j.dsr2.2008.05.006

Wilson, S. E., D. K. Steinberg, and K. O. Buesseler. 2008. Changes in fecal pellet characteristics with depth as indicators of zooplankton repackaging of particles in the mesopelagic zone of the subtropical and subarctic North Pacific Ocean. Deep Sea Res 2 Top Stud Oceanogr **55**: 1636–1647. doi:10.1016/j.dsr2.2008.04.019

Youngbluth, M. J., T. G. Bailey, P. J. Davoll, C. A. Jacoby, P. I. Blades-Eckelbarger, and C. A. Griswold. 1989. Fecal pellet production and diel migratory behavior by the euphausiid Meganyctiphanes norvegica effect benthic-pelagic coupling. Deep Sea Research Part A, Oceanographic Research Papers **36**: 1491–1501. doi:10.1016/0198-0149(89)90053-8
